# Dengue virus infection in *Aedes aegypti* mosquito brains elicits minimal transcriptional response

**DOI:** 10.64898/2026.06.10.731349

**Authors:** Umberto Palatini, Stéphanie Dabo, Adriana Rosa-Villegas, Yael N. Tsitohay, Alexandra E. DeFoe, Nadav Shai, Louis Lambrechts, Leslie B. Vosshall

## Abstract

Billions of people each year are at risk from infection by dengue, Zika, yellow fever, and chikungunya viruses, which are transmitted by female Aedes aegypti mosquitoes. Mosquitoes themselves are infected by these arboviruses, but how the mosquito nervous system responds to arboviral infection is unknown. We combined whole-mount immunofluorescence with single-head bulk RNA-sequencing to characterize dengue virus (DENV) infection in the brain of Aedes aegypti. DENV productively infects brain cells in a bimodal pattern: individual brains showed either sparse or widespread infection, with no intermediate phenotypes. An infectious blood meal altered thousands of genes, including 64 immunity genes, at 7 days post-feeding (DPF), yet active viral replication in the head did not increase the transcriptional response. Heads with and without detectable DENV showed minimal transcriptional differences, with no induction of canonical immune effectors. Despite productive infection, the mosquito brain tolerates DENV replication with minimal transcriptional response.

---

Arthropod-borne viruses (arboviruses) represent a major and growing threat to global public health. Dengue virus (DENV) alone is estimated to cause more than 390 million infections annually, with incidence expected to rise as climate change extends the range of mosquito vectors^1,2^. The primary arbovirus vector in Africa and the Americas is *Aedes aegypti*, a highly anthropophilic mosquito whose females require blood meals to produce eggs and can feed on multiple humans during their 2-6 week lifespan^3–6^. *Aedes aegypti* acquire arboviruses by feeding on viremic human hosts. The virus then replicates sequentially in the midgut, disseminates through the hemocoel, and reaches the salivary glands, enabling transmission during subsequent blood meals^7^.

The arbovirus replication cycle depends on the ability of mosquitoes to tolerate high levels of viral replication in most tissues without succumbing to infection^7,8^. Disease tolerance, the ability to limit the fitness costs of infection without eliminating the pathogen, and resistance jointly shape vectorial capacity: tolerance by promoting mosquito survival, resistance by reducing vector competence^9,10^. Tolerance likely reflects adaptation on both sides: mosquitoes have evolved mechanisms to limit the severity of viral infection, while arbovirus virulence in insect vectors is shaped by the trade-off between within-host replication and the preservation of mosquito survival and behavior needed for viral transmission^11–13^. Arboviral resistance in vector mosquitoes has been thoroughly explored, but we are only beginning to understand tolerance mechanisms^10,14,15^, with recent work implicating compensatory tissue repair^16^, virus-derived DNA^17,18^, RNA interference^19–21^, protective factors^22^, and the microbiota^23^. How different tissues achieve tolerance remains poorly characterized.

Surviving infection is only the first requirement for transmission. For both pathogen and vector to succeed, infected vectors must maintain the sensory and motor functions needed to find vertebrate hosts and feed on their blood. Some pathogens do more, actively modifying vector behavior to enhance their own transmission. *Leishmania* parasites modify sand fly feeding behavior and physiology to enhance transmission^24,25^. *Plasmodium*-infected *Anopheles* mosquitoes show stage-dependent behavioral changes: reduced feeding motivation during non-transmissible oocyst stages and enhanced human host-seeking when carrying transmissible sporozoites^26–28^. This pattern suggests adaptive manipulation benefiting the parasite, and some insect viruses employ similar strategies: baculoviruses induce tree-top disease in caterpillars, driving infected animals to climb to elevated positions before death to maximize viral dispersal^29^. These examples suggest that arboviruses could similarly enhance their own transmission by modifying mosquito host-seeking or biting behavior.

The literature describing the effect of arboviral infection on mosquito behavior remains contradictory and inconclusive^30^. Studies in different mosquito species examining dengue, Zika, La Crosse, and West Nile viruses have reported inconsistent effects: West Nile and La Crosse viruses appeared to reduce host-seeking activity^31,32^, while DENV increased antennal sensitivity to human odors^33^. Studies that examined the ability of the mosquito to feed on blood show similar inconsistency. Some found no effect of DENV infection on probing behavior, while others reported subtle changes^34–38^. DENV infection also alters the skin microbiota of infected human and rodent hosts, increasing their attractiveness to mosquitoes^39^. Whether any of these effects reflect direct viral manipulation or general pathophysiology remains unknown, in part because of heterogeneity in virus strains and vector species, small sample sizes, and lack of standardized methodology^30^. Recent work indicates that inconsistent reports in the literature may in fact reflect true biological differences among mosquito-pathogen pairs^38^.

The nervous system represents an obvious target for behavioral manipulation by the virus, yet arboviral infection of mosquito brains remains almost entirely uncharacterized. DENV antigens have been detected in mosquito neural tissue by immunofluorescence of head squashes^40^ and in cultured neurons, where the neural protective factor *Hikaru genki* (*AAEL004725*) was observed to limit virus-induced neuronal death^22,41^. These observations confirm that orthoflaviviruses can reach and replicate in the nervous system, but neither the spatial distribution of infection within the brain nor the transcriptional consequences have been examined. This contrasts with the well-studied biology of midgut and salivary glands, where arbovirus infection induces hundreds of differentially expressed genes, including canonical immune genes^36,42,43^. Whether the brain mounts similar responses or remains transcriptionally quiescent is unknown. Interpreting any brain transcriptional response requires an additional distinction: ingesting virus-containing blood could alter brain gene expression through circulating viral antigens or systemic immune signals, independent of the virus establishing local infection in neural tissue. Distinguishing transcriptional changes driven by viral exposure from those induced by active replication in the brain is essential for determining whether the brain mounts tissue-specific defenses or tolerates viral replication with minimal transcriptional consequence.

Although neurological complications of dengue are rare in humans, DENV can cross the blood-brain barrier and infect neurons, astrocytes, microglia, and endothelial cells of the central nervous system^44–46^. The clinical consequences can be severe, including seizures, reduced consciousness, and death^46,47^. In contrast, *Aedes aegypti* mosquitoes tolerate productive viral replication across most tissues yet survive and continue to host-seek and blood-feed^10,14,22^. What molecular features of the mosquito brain allow it to tolerate a virus that can cause lethal neuropathology in its vertebrate host is unknown.

Here we characterize the spatial distribution and transcriptional consequences of DENV infection in the *Aedes aegypti* female brain. Using single-head bulk RNA-sequencing (RNA-seq) paired with reverse transcription quantitative polymerase chain reaction (RT-qPCR) validation, we distinguish transcriptional responses to viral exposure from responses to active DENV brain infection. Using immunofluorescence, we demonstrate the presence of DENV virions in the brain with a bimodal distribution: brains showed either minimal or widespread viral antigen, with no intermediate phenotypes. Despite extensive infection in some individuals, oral exposure to live DENV did not increase mortality, and DENV-exposed mosquitoes showed intact gross motor responses prior to dissection. Exposure to live DENV drove widespread transcriptional changes in thousands of genes, but comparing DENV-positive to DENV-negative heads revealed only five upregulated genes at 7 DPF under a criterion that included carcass infection, and none at later time points. This minimal response contrasts with other tissues and suggests that the brain represents a uniquely tolerant organ.

## RESULTS

### Experimental infection of *Aedes aegypti* female mosquitoes with dengue virus

To determine whether dengue virus (DENV) reaches the mosquito brain, we orally exposed female *Aedes aegypti* mosquitoes (Liverpool strain)^48^ to DENV (serotype 1)^49^. Female mosquitoes were offered artificial blood meals containing either live DENV [10⁶ focus-forming units (FFU)/mL; n=505 animals], heat-inactivated DENV (n=214), to control for the presence of viral proteins without active replication^50^, or no virus (n=204), to control for blood-feeding-induced physiological changes (**Figure 1A**). All mosquitoes used in downstream analyses were fed on the same day and time to minimize temporal variation. Surviving animals were dissected in the morning at 7, 11, and 14 days post-feeding (DPF). At dissection, heads, legs, and carcasses (body minus head and legs) were processed separately for downstream RT-qPCR and either RNA-seq or immunofluorescence staining (see Methods).

**Figure 1.**
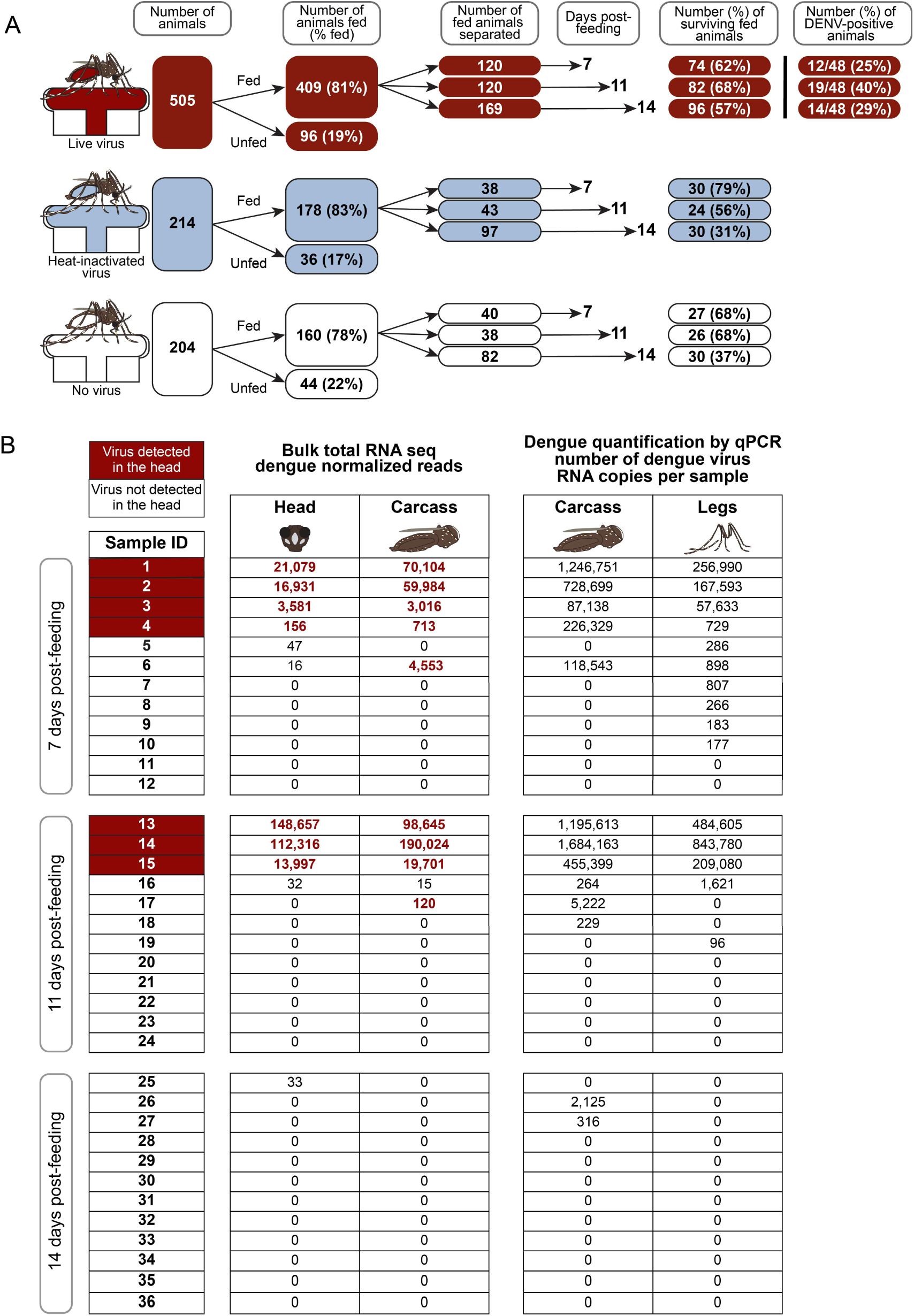
| DENV is detected in a subset of orally exposed *Aedes aegypti* mosquitoes. (**A**) Experimental workflow and sample sizes. Female mosquitoes were offered blood meals containing live DENV (dark red), heat-inactivated virus (blue), or no virus (white). Fed mosquitoes were separated into three cohorts and dissected at 7, 11, and 14 DPF. Values in rectangles indicate the number of individuals that progressed to the next step. DENV-positive individuals at each time point are at the top right. (B) DENV detection in individual mosquitoes by RNA-seq and RT-qPCR. Left, normalized DENV-assigned reads from total RNA-seq of single heads and paired carcasses. Right, total DENV genomic RNA copies quantified by RT-qPCR in paired carcasses and pooled legs from the same individuals. Sample IDs shaded in dark red at the left denote mosquitoes with DENV detected in the head (>100 DENV genomic reads in the head RNA-seq library). The normalized reads of RNA-seq samples with >100 DENV genomic reads in either head or carcass are indicated with dark red text. See also **Tables S1 and S2**.

Feeding rates were comparable across the three groups (live virus, 81%; heat-inactivated, 83%; no virus, 78%) (**Figure 1A, Table S1**), indicating that viral material did not impact blood meal palatability. Survival to the time of dissection was 62% in the live-virus group, 47% in heat-inactivated, and 52% in no-virus controls (**Figure 1A, Table S1**). These rates are lower than those typically observed in standard insectary conditions and reflect the constraints of blood-feeding experimental logistics in an Arthropod Containment Level 3 (ACL-3) laboratory, including cold-anesthesia, sorting on ice, increased housing density, and discontinuous access to fresh sucrose, all of which reduce *Aedes aegypti* adult survival^51,52^. Survival did not differ significantly among experimental groups at 7 or 11 DPF (Chi-square test, p = 0.14 and p = 0.31, respectively). At 14 DPF, survival was higher in the live virus group than in heat-inactivated or no-virus controls (Fisher’s exact test, p < 0.001 and p = 0.003, respectively), most likely reflecting differential housing density from unequal starting group sizes rather than a protective effect of virus exposure (**Figure 1A, Table S1**).

Of the 419 surviving and dissected mosquitoes, 311, including 144 DENV-exposed heads (48 per time point) and all surviving controls (84 heat-inactivated, 83 no-virus), were shipped from Institut Pasteur to The Rockefeller University. The remaining 108 mosquitoes were stored at the Institut Pasteur as backup samples.

We assessed DENV infection in the carcass and legs of live virus-exposed mosquitoes by probe-based RT-qPCR targeting the *NS5* viral gene, normalized against two *Aedes aegypti* housekeeping genes: *Actin5c* (*AAEL011197*) and *RPS17* (*AAEL004175*)^53^ (**Table S2**). Carcass infection rates did not differ significantly over time (chi-square test, p=0.28), with viral RNA detected in 25.0% [12/48; 95% Confidence Interval (CI): 14.9–38.8%], 39.6% (19/48; 95% CI: 27.0–53.7%), and 29.2% (14/48; 95% CI: 18.2–43.2%) of mosquitoes at 7, 11, and 14 DPF, respectively. Viral loads spanned more than three orders of magnitude at each time point (**Figure 1B, Table S2**), consistent with the high inter-individual variability documented for oral arbovirus infection in *Aedes aegypti*^43,54–56^. From our cohort of mosquitoes, 12 DENV-exposed heads per time point (n=36 total) were selected for bulk RNA-seq based on RNA integrity, alongside 7-8 heads per time point each for heat-inactivated and no-virus controls (n=47 in total). The remaining heads were processed for whole-mount immunofluorescence; 58 DENV-exposed brains and 16 brains from each control condition were successfully dissected, stained, and imaged on a confocal microscope (**Tables S1 and S2**).

We defined RNA-seq samples (heads or carcasses) as “virus-positive” if they contained more than 100 genomic reads mapping with high confidence to the genome sequence of our DENV isolate, a conservative threshold chosen to exclude stochastic read mismapping while retaining low-level infections. Among the live virus-exposed heads used for total RNA-seq, DENV genomic reads were detectable in 4 of 12 at 7 DPF, 3 of 12 at 11 DPF, and 0 of 12 at 14 DPF (**Figure 1B**).

All head-positive mosquitoes had robust viral reads in their paired carcasses, whereas two individuals, one at 7 and one at 11 DPF, were DENV-positive in the carcass but not the head (**Figure 1B, Table S2**). None of the 12 mosquitoes sequenced at 14 DPF had detectable DENV reads in the head.

However, head dissemination rates did not differ significantly across time points (4/12, 3/12, and 0/12 at 7, 11, and 14 DPF; Fisher’s exact test, all pairwise p > 0.09), brain infection prevalence by immunofluorescence was likewise stable (3/19, 8/19, and 5/20 infected brains; all pairwise p > 0.15; **Figure 2**), and systemic infection rates in carcasses and legs were similarly stable across time points (**Table S2**). The absence of DENV reads in heads at 14 DPF therefore likely reflects low prevalence in a small subset of sequenced heads rather than viral clearance. Among RT-qPCR-positive carcasses, viral loads spanned four orders of magnitude, and within RNA-seq-positive heads, normalized DENV read counts ranged from 156 to 148,657, mirroring the variation seen in the full RT-qPCR cohort and consistent with the known heterogeneity in susceptibility and antiviral responses, as documented in both vertebrate hosts^57^ and mosquito vectors^55,56,58–61^.

**Figure 2.**
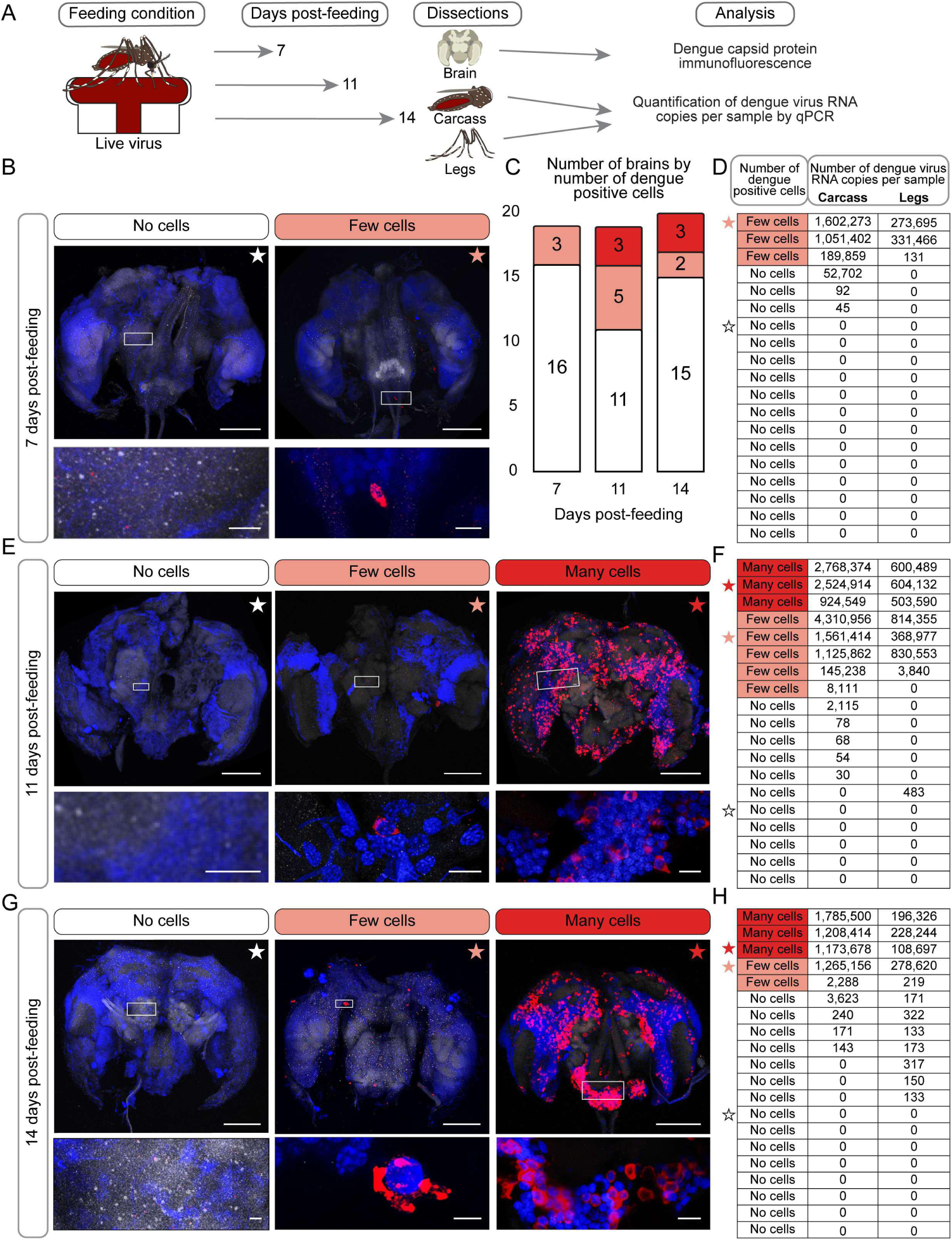
| DENV productively infects *Aedes aegypti* brains in a bimodal pattern. **(A)** Experimental workflow. DENV-exposed mosquitoes were dissected at 7, 11, and 14 DPF. Brains were processed for immunofluorescence; carcasses and legs were processed for RT-qPCR. **(B, E, G)** Representative maximum-intensity projections of whole-mount mosquito brains at 7 **(B)**, 11 **(E)**, and 14 **(G)** DPF, categorized by the number of cells showing DENV complex antigen staining: no cells, few cells, or many cells. DENV complex antigen, magenta; N-cadherin, grey; DAPI, blue. Scale bars, 100 μm (main images), 10 μm (all insets except 14DPF “few cells”, which has a 5 μm scale bar). Insets show enlarged views of the boxed regions. Stars link each representative brain to the corresponding individual’s viral load data in **(D, F, H)**. **(C)** Number of brains by infection category across time points. **(D, F, H)** DENV RNA copy numbers quantified by RT-qPCR in paired carcass and leg samples for each individual brain imaged at 7 (n=19) **(D)**, 11 (n=19) **(F)**, and 14 (n=20) **(H)** DPF. Rows are sorted by infection category and viral load. See also **Figure S1** and **Tables S1 and S2**.

### DENV productively infects mosquito brains

To determine whether DENV productively infects the brain, we performed whole-mount immunofluorescence on brains dissected from paraformaldehyde-fixed heads of live DENV-exposed mosquitoes at 7, 11, and 14 DPF (**Figure 2A**). We used an antibody against the DENV complex antigen to detect intracellular viral antigen, providing direct evidence of virion presence within cells. Brains were co-stained with an anti-N-cadherin antibody to visualize neuropil structure and DAPI to label nuclei and imaged by confocal microscopy. Of 72 DENV-exposed heads processed for immunofluorescence (24 per time point), 58 brains were successfully dissected, stained, and imaged: 19 at 7 DPF, 19 at 11 DPF, and 20 at 14 DPF. We also imaged 16 brains from heat-inactivated and 16 from no-virus controls as negative controls (**Figure S1**, **Tables S1 and S2**).

At 7 DPF, 16 of 19 brains (84.2%) showed no detectable viral antigen (”no cells”), and three brains (15.8%) contained fewer than 15 visible DENV-positive cells, exhibiting the characteristic perinuclear localization of viral particles^62,63^ (**Figure 2B-D**). We defined these brains as having “few cells”. By 11 DPF, a bimodal pattern emerged: brains harbored either scattered DENV-positive cells (“few cells”) or widespread antigen spanning multiple regions (“many cells”), with no intermediate phenotypes. Of 19 brains imaged at 11 DPF, 11 (57.9%) showed no DENV-positive cells, 5 (26.3%) showed few DENV-positive cells, and 3 (15.8%) showed many DENV-positive cells (**Figure 2C, E**-**F**). This bimodal distribution persisted at 14 DPF, where 15 of 20 brains (75.0%) showed no DENV-positive cells, 2 (10.0%) showed few DENV-positive cells, and 3 (15.0%) showed many DENV-positive cells (**Figure 2C, G-H**).

Brains classified as “many cells” showed viral antigen distributed across multiple brain regions. No DENV-positive cells were detected in brains from heat-inactivated virus or no-virus control mosquitoes at any time point examined (Figure S1), confirming that the DENV complex antigen staining reflects productive intracellular infection rather than passive antigen accumulation or nonspecific antibody binding. Our data provide spatially resolved characterization of DENV infection in the *Aedes aegypti* brain.

### Single-head RNA sequencing reveals transcriptional changes in DENV-exposed brains

We first performed short-read RNA-seq on poly(A)-enriched mRNA from pools of 10 mosquito heads (with appendages removed) collected at 14 DPF to ask whether we could detect transcriptional changes in DENV-exposed brains. For this experiment only, we sequenced the antennal transcriptomes as well, pooling the 20 antennae derived from the heads of each 10-mosquito biological replicate. We compared four pools from DENV-exposed mosquitoes matched for viral titer (as measured by RT-qPCR; **Table S3**) against four pools from no-virus controls (**Figure S2A**, **Data S1 and S2**). To identify differentially expressed genes (DEGs), we performed pairwise comparisons between experimental groups using DESeq2^64^.

This analysis identified only 7 DEGs in heads and 11 in antennae (**Figure S2B, Data S3 and S4**), far fewer than the hundreds of DEGs reported in DENV-infected midgut and salivary glands when compared to uninfected tissues^13,16,36,43^.

Because data from pools are an average across individuals with variable infection status and response, we could not determine whether these minimal differences reflected biological signal or the averaging of heterogeneous individual responses within pools. A previous study using pooled *Aedes aegypti* antennae similarly found that DENV-1 infection altered some neural signaling transcripts and few chemosensory genes but led to minimal global transcriptional changes^33^, consistent with our pooled results (**Figure S2B**). These initial results motivated a revised experimental design using single-mosquito head bulk RNA-seq of total rRNA-depleted RNA libraries, which captures both mosquito transcripts and DENV reads, paired with matched probe-based RT-qPCR validation from each individual sequenced. This approach allowed us to identify transcriptional changes specifically associated with active infection rather than viral exposure alone, at the level of individual mosquitoes rather than pools.

We performed bulk RNA-seq on 83 individual heads with head appendages removed from all three experimental conditions (36 live virus, 24 heat-inactivated, 23 no virus) and sequenced the corresponding carcasses belonging to the same live virus-exposed mosquitoes (12 per time point; 36 total; **Data S5 and S6**).

Principal component analysis of global gene expression at 7 DPF revealed separation between live-virus-exposed samples and controls along PC1 (40% of variance) and PC2 (7.5%), with DENV-exposed heads occupying distinct transcriptional space regardless of whether DENV reads were detected in the head (Figure 3A). A heatmap of z-score-normalized expression across all 13,272 expressed genes at 7 DPF confirmed this pattern: live-virus-exposed heads clustered together irrespective of head viral read detection, while heat-inactivated and no-virus controls formed a separate cluster (**Figure S3**). By 11 and 14 DPF, samples from all experimental conditions largely overlapped in PCA space (**Figure 3B and 3C**). At 7 DPF, exposure to live DENV, rather than detectable head infection, accounted for the observed transcriptional variation, and this effect was no longer apparent at later time points.

Using the same DESeq2 analysis applied to pools (padj < 0.01, |log₂FC| ≥ 1), we identified similar numbers of DEGs at 7 DPF in both virus-positive heads (3,206 DEGs) and virus-negative heads (3,398 DEGs) compared to no-virus controls, reinforcing that exposure to live virus rather than detectable head infection drove transcriptional changes at this time point (**Figure 3D, Table S4, Data S7**). Fewer DEGs emerged when comparing virus-positive (725 DEGs) and virus-negative (476 DEGs) heads against heat-inactivated controls, suggesting partial transcriptional overlap between heat-inactivated and live virus exposure. By 11 DPF, DEG counts between DENV-exposed and control groups fell sharply (19–45 DEGs across comparisons), and by 14 DPF, virus-negative heads showed 81 DEGs versus no-virus controls and 135 versus heat-inactivated controls (**Figure 3D, Data S7**), consistent with the convergence observed by PCA. The infectious blood-meal–driven response was therefore transient, peaking at 7 DPF and largely resolving by 11 DPF. Despite clear evidence of viral replication in a subset of brains, including widespread infection observed by immunofluorescence in other individuals from the same cohort (**Figure 2**), heads with confirmed DENV reads showed zero differentially expressed genes compared to virus-negative heads at any time point (**Figure 3D**).

**Figure 3.**
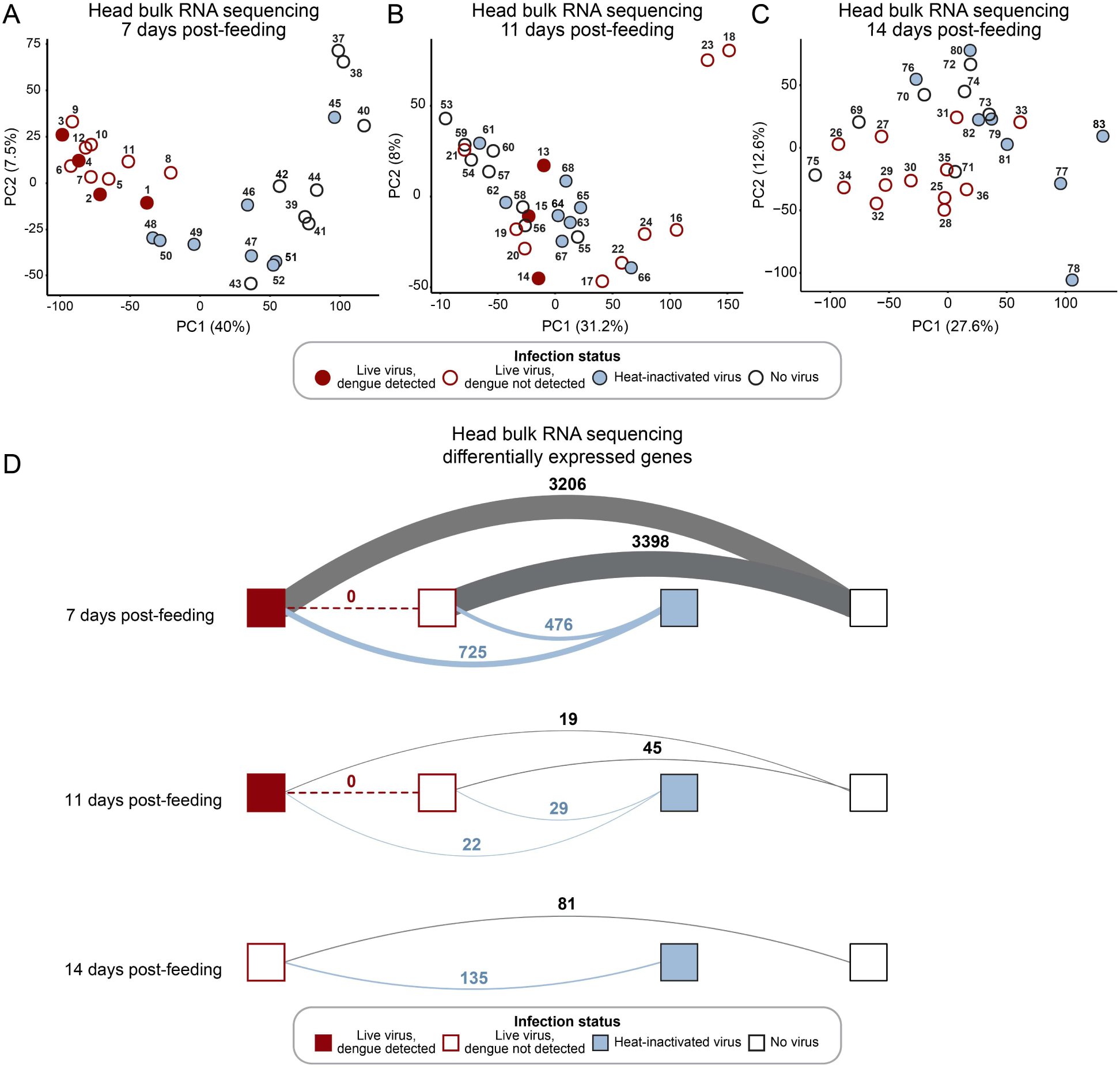
| DENV exposure, and not infection, drives transcriptional changes at 7DPF. **(A-C)** Principal component analysis (PCA) of gene expression in individual mosquito heads at 7 **(A)**, 11 **(B)**, and 14 **(C)** DPF. Each point represents one individual, labeled by sample ID. The percentage of variance explained by each principal component is indicated on axes. **(D)** Number of significantly differentially expressed genes (padj < 0.01, |log₂FC| ≥ 1) identified by pairwise comparisons between experimental groups at each time point. Boxes represent experimental conditions; connecting bands indicate pairwise comparisons, with the number of differentially expressed genes shown. The dashed red band and circled value highlight the comparison between virus-positive and virus-negative samples within live virus-exposed mosquitoes (virus-positive: > 100 viral reads in head RNA-seq; n=4 vs. n=8 at 7 DPF; n=3 vs. n=9 at 11 DPF, n=0 vs. n=12 at 14 DPF). For all panels: white fill, no-virus control; blue fill, heat-inactivated virus; dark red outline with white fill, live virus exposure, virus-negative; solid dark red fill, live virus exposure, virus-positive. See also **Figure S3** and **Data S7**.

### Virus exposure, rather than infection in the head, drives transcriptional changes in immunity and brain-associated genes

To further characterize the transcriptional changes detected at 7 DPF, we examined two curated gene sets: 413 putative immunity-related genes spanning 27 functional categories (including antimicrobial peptides, RNA interference components, and Toll, IMD, and JAK-STAT pathway genes), and 48 high-confidence brain-associated genes (**Data S8**). These gene sets were curated from available datasets and current annotations of mosquito immunity and neural functions^65,66^. Compared to no-virus controls, virus-positive heads showed 64 differentially expressed immunity genes and 18 brain-associated marker genes, while virus-negative heads showed 68 and 15, respectively (**Table S5, Data S8**). The most responsive immunity categories included the IMD pathway (5/12 genes, 42%), autophagy (6/19 genes, 32%), and small RNA regulatory pathways (11/37 genes, 30%). Antimicrobial peptides, prophenoloxidases, β-1,3-glucan binding proteins, and the NF-κB transcription factor *Relish* showed no differential expression. Among brain-associated markers, glutamate and serotonin transporters and the biogenic amine biosynthetic enzymes tyramine β-hydroxylase and tyrosine decarboxylase were upregulated, while the short neuropeptide F precursor and myosuppressin receptor were downregulated. The neural protective factor *Hikaru genki* (*AAEL004725*) was upregulated ∼3-fold in both DENV-exposed groups relative to no-virus controls, confirming earlier observations^22^.

Heat-inactivated virus elicited a partial response (**Table S5, Data S8**). Compared to heat-inactivated controls, virus-positive and virus-negative heads contained only 14 and 16 immunity DEGs, respectively, versus 64 and 68 DEGs against no-virus controls, indicating that 50 of 64 (78%) of the immunity DEGs induced by live virus are also triggered by heat-inactivated virus. For brain-associated markers, the overlap was nearly complete: 17 of 18 (94%) of the genes differentially expressed in virus-positive heads versus no-virus were no longer detected when compared against heat-inactivated controls. Although heat-inactivated and no-virus heads showed broadly similar global transcriptional profiles (**Figure 3A-C, Figure S3**), this targeted analysis distinguishes them: heat-stable viral components, likely complex antigen or other structural proteins, are sufficient to drive most of the immunity and brain-associated transcriptional response we detect, while the residual signal in live-exposed heads may require replication intermediates or heat-labile viral structures.

### Head infection alone yields zero differentially expressed genes; carcass infection reveals five

DENV genomic reads and viral antigen were present in a subset of brains (**Figure 1, Table S2**), yet differential expression analysis comparing virus-positive and virus-negative heads (padj < 0.01, |log₂FC| ≥ 1) identified zero differentially expressed genes at any time point (**Figure 3D**).

DENV can replicate in the carcass without disseminating to the head, so we asked whether infection in the thorax and abdomen altered head gene expression independent of head infection. We expanded the virus-positive criterion to include any mosquito with >100 viral reads in either the head or the paired carcass library. This reclassified one additional mosquito at 7 DPF (n=5 virus-positive vs. n=7 virus-negative) and one additional mosquito at 11 DPF (n=4 vs. n=8). All differential expression analyses were performed on the head transcriptome; carcass RNA-seq was used only to assign group membership.

Under this broader criterion we identified five upregulated genes at 7 DPF and no differentially expressed genes at 11 or 14 DPF (**Table 1, Table S4, Figure 4, Figure S3**). These genes encode Histone H4 (*AAEL013709*), thioredoxin peroxidase (*AAEL013528*), ribulose-5-phosphate-3-epimerase (*AAEL010505*), 40S ribosomal protein S15 (*AAEL011656*), and a PBAN/pyrokinin neuropeptide precursor (*AAEL012060*), with log₂ fold-changes between 1.03 and 1.17 and adjusted p-values from 9.7×10^-8^ to 1.3×10^-4^. Under the stricter head-only criterion (n=4 vs. n=8), four of these five genes retained padj < 0.01 but their fold-changes fell below the |log₂FC| ≥ 1 cutoff. *AAEL012060* and *AAEL013709* narrowly missed the cutoff (log₂FC = 0.989 and 0.997; padj = 1.6×10^-4^ and 2.8×10^-4^), and *AAEL011656* (40S ribosomal protein S15) did not reach significance (padj = 7.0×10^-2^) (**Table 1**).

**Figure 4.**
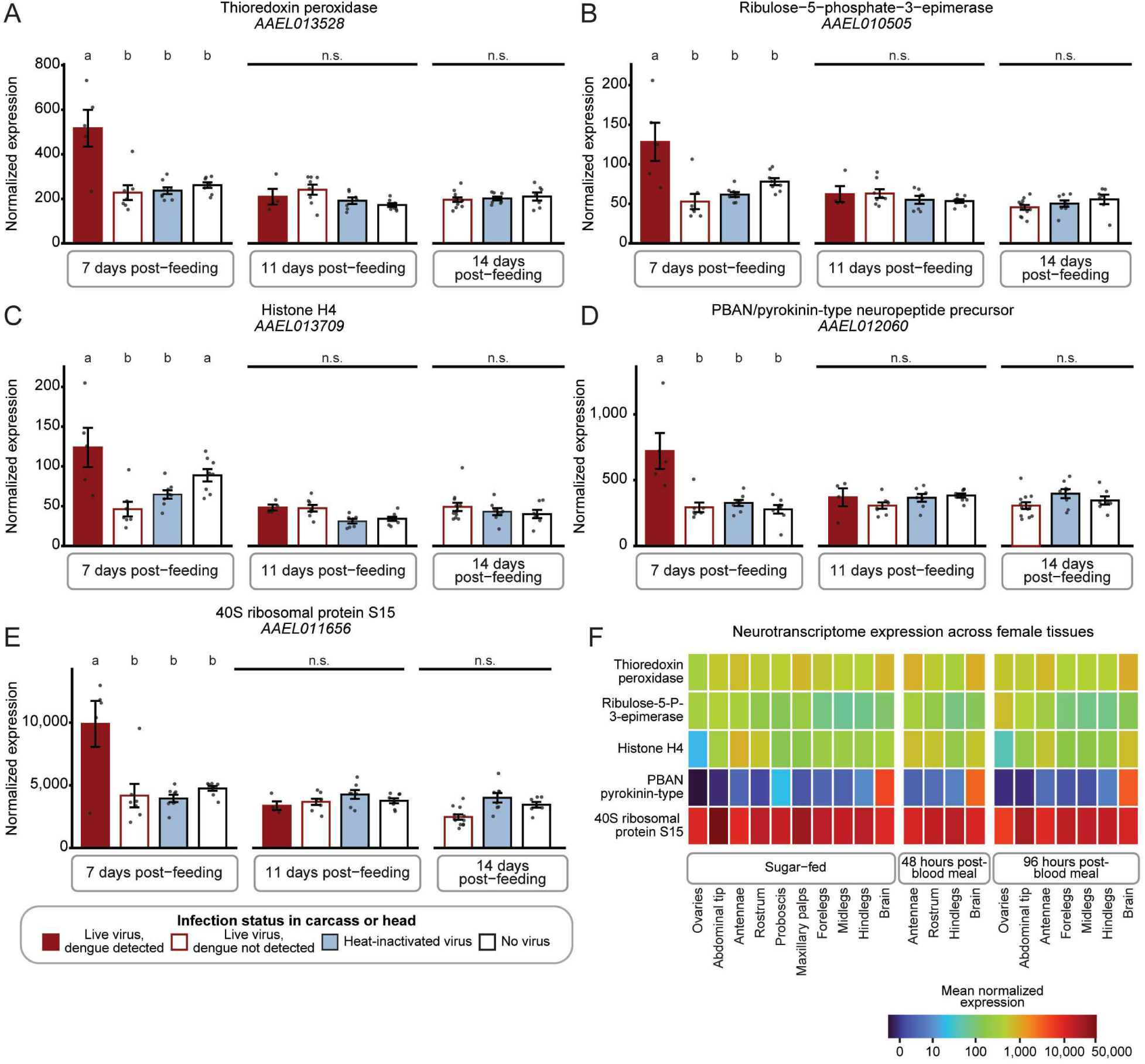
| Five genes are upregulated at 7 DPF in heads of mosquitoes that are virus-positive in the head or carcass. (**A–E**) Normalized RNA-seq counts at 7, 11, and 14 DPF for the five genes upregulated in virus-positive heads at 7 DPF: thioredoxin peroxidase (*AAEL013528*, **A**), ribulose-5-phosphate-3-epimerase (*AAEL010505*, **B**), Histone H4 (*AAEL013709*, **C**), PBAN/pyrokinin-type neuropeptide precursor (*AAEL012060*, **D**), and 40S ribosomal protein S15 (*AAEL011656*, **E**). Bars indicate mean ± SD with individual data points overlaid. Conditions are color-coded: white, no virus (n=8 at 7 and 11 DPF; n=7 at 14 DPF); blue, heat-inactivated DENV (n=8 per time point); white with dark red outline, live DENV exposure with virus not detected (n=7 at 7 DPF; n=8 at 11 DPF); solid dark red, live DENV exposure with virus detected (n=5 at 7 DPF; n=4 at 11 DPF). Differential expression was assessed by Wald test in DESeq2^64^ with Benjamini-Hochberg correction; significance threshold padj < 0.01. Letters above bars denote significant pairwise differences within each time point; groups sharing a letter are not significantly different. n.s., not significant. **(F)** Expression of the five upregulated genes across the indicated female *Aedes aegypti* tissues, in three physiological states: sugar-fed, 48 h post-blood meal, and 96 h post-blood meal. Data are from the neurotranscriptome dataset^67^. Color scale represents mean normalized expression on a log scale. See also **Figure S3, Data S6 and S10**.

**Table 1.** | Five DENV-responsive genes identified by differential expression analysis of mosquito heads at 7 DPF. Differentially expressed genes (padj < 0.01, |log₂FC| ≥ 1) were called from RNA-seq of single mosquito heads, with samples grouped under two criteria: by head viral RNA alone (By Head) or by head or carcass viral RNA (By Carcass/Head). Both criteria were applied to the same head transcriptome data; carcass data were used only for group assignment. Rows are sorted by adjusted p-value within the head-or-carcass criterion. Fold-changes [log2(FC)] and adjusted p-values (padj) were calculated by DESeq2^64^ (Wald test, Benjamini-Hochberg correction). The mean expression level of each gene is indicated in the “baseMean” column. Full DESeq2 output for all DEGs (padj < 0.01, |log₂FC| ≥ 1) is provided in **Data S7 and S9**.

| Comparison vs. virus-negative | Gene ID | baseMean | By Carcass/Head |  | By Head only |  |
| --- | --- | --- | --- | --- | --- | --- |
|  |  |  | log<sub>2</sub>(FC) | p<sub>adj</sub> | log<sub>2</sub>(FC) | p<sub>adj</sub> |
| 7 DPF | AAEL013528 | 232.2 | 1.070 | 9.7×10<sup>-8</sup> | 0.759 | 1.5×10<sup>-3</sup> |
|  | AAEL010505 | 61.2 | 1.100 | 8.0×10<sup>-6</sup> | 0.654 | 8.8×10<sup>-3</sup> |
|  | AAEL013709 | 54.1 | 1.171 | 3.5×10<sup>-5</sup> | 0.997 | 2.8×10<sup>-4</sup> |
|  | AAEL012060 | 360.8 | 1.087 | 5.9×10<sup>-5</sup> | 0.989 | 1.6×10<sup>-4</sup> |
|  | AAEL011656 | 4094.4 | 1.031 | 1.3×10<sup>-4</sup> | 0.333 | 7.0×10<sup>-2</sup> |

Expression of these five genes was elevated in virus-positive samples at 7 DPF and returned to baseline by 11 DPF, identifying the infection response as transient (**Figure 4A-4E).** We could not test the comparison at 14 DPF because no virus-positive samples were obtained at that time point. At all three time points, expression in virus-negative samples was indistinguishable from the no-virus and heat-inactivated-virus control groups, indicating that upregulation tracks productive infection rather than blood-meal exposure to viral particles (**Figure 4A-E**).

One additional gene, *AAEL021837* (annotated as an RRM domain-containing protein, UniProt ID: Q16H40), was initially called differentially expressed at 11 DPF under the broader criterion. *AAEL021837* and its paralog *AAEL002980* share 100% nucleotide identity over 930 bp and the same UniProt entry, and their expression is mutually exclusive across all 83 samples (Pearson r = −0.90) (**Figure S4**). This pattern is consistent with read-assignment ambiguity between identical loci, possibly arising from a genome misassembly, and is therefore a technical artifact. We therefore excluded *AAEL021837* from the DEG set. No other DEG in the virus-positive versus virus-negative comparison showed this paralog-aliasing pattern.

To determine whether the five upregulated genes are expressed broadly or show tissue specificity, we examined their expression across uninfected female *Aedes aegypti* tissues using our published neurotranscriptome dataset (**Figure 4F**). Histone H4, thioredoxin peroxidase, ribulose-5-phosphate-3-epimerase, and 40S ribosomal protein S15 were expressed at moderate to high levels across all tissues, consistent with their housekeeping or metabolic functions. *AAEL012060*, the PBAN/pyrokinin neuropeptide precursor, was enriched in the brain relative to antennae, rostrum, proboscis, maxillary palps, legs, abdominal tip, and ovaries, and this brain enrichment was consistent across sugar-fed, 48-hour post-blood meal, and 96-hour post-blood meal states (**Figure 4F, Data S10**).

No canonical immune gene (pattern recognition receptors, antimicrobial peptides, RNA interference machinery, and core signaling components of the Toll, IMD, and JAK-STAT pathways) was differentially expressed in the virus-positive versus virus-negative comparison at any time point, under either grouping criterion (**Data S8-S9**). The only immunity-classified gene among the five was thioredoxin peroxidase (*AAEL013528*), which functions in oxidative stress management rather than canonical antiviral defense.

## DISCUSSION

Dengue, Zika, yellow fever, and chikungunya viruses together threaten more than five billion people each year^2^. How these arboviruses interact with the *Aedes aegypti* nervous system has remained largely uncharacterized, despite the central role of mosquito behavior in transmission. Here we provide the first spatial and transcriptional characterization of dengue virus (DENV) infection in the *Aedes aegypti* female brain combining whole-mount immunofluorescent staining of DENV complex antigen and single-head transcriptomic profiling. DENV productively infects the brain, exposure to virus through an infected blood meal elicits a measurable transcriptional response in the head, however active replication adds almost nothing to that response.

### DENV establishes productive infection in the mosquito brain

DENV infects *Aedes aegypti* brain cells in a bimodal pattern: Of 58 brains examined, 16 contained DENV antigen; of these, 10 showed only a few scattered positive cells and 6 showed widespread antigen across multiple brain regions, with no intermediate phenotypes. This bimodal distribution was consistent across time points (Fisher’s exact test, all pairwise p > 0.15). Earlier studies detected DENV antigen in squashed mosquito head tissue but lacked the spatial resolution to define where, and how much, the virus replicates^40^. Our whole-mount immunofluorescence localizes infection to discrete cells within the brain and shows that infection is not a graded continuum.

The biological basis of the bimodal distribution is unclear. The two outcomes may reflect stable inter-individual differences in permissiveness, with some mosquitoes restricting viral spread within the brain and others permitting widespread replication. Alternatively, it may represent successive stages of the same infection dynamics, with “few cells” preceding “many cells” as infection expands. Cross-sectional sampling cannot separate these models; resolving them requires live imaging of individual mosquitoes over time.

### The mosquito brain mounts a minimal transcriptional response to DENV infection

Because each head was sequenced individually, we could distinguish transcriptional changes driven by ingestion of an infectious blood meal from those driven by active viral replication in the head and/or carcass. Globally, live DENV-exposed heads showed a transcriptional profile distinct from both heat-inactivated and no-virus heads, with thousands of differentially expressed genes distinguishing each pairwise comparison at 7 DPF. Within the curated immunity gene set, however, the overlap between live-virus and heat-inactivated responses was substantial: 78% of the immunity-gene response (52 of 64 DEGs at 7 DPF) and nearly all of the brain-marker response (17 of 18 DEGs) seen against no-virus controls were also induced by heat-inactivated virus. Heat-stable viral components are therefore sufficient to drive most of the immunity and brain-associated transcriptional response at 7 DPF, even though the broader transcriptome clearly distinguishes the two conditions.

Under the strict head-only criterion, no genes were differentially expressed between virus-positive and virus-negative heads at any time point. Under the broader criterion defining infection by DENV detection anywhere in the mosquito, only five genes reached significance, all upregulated at 7 DPF. The five upregulated genes encode proteins consistent with a generic cellular accommodation of viral replication rather than a dedicated antiviral program. Histone H4 (*AAEL013709*) is a core nucleosome component whose upregulation may reflect increased chromatin demand or a more direct role in infection, as histone H4 has been reported to act as a DENV proviral host factor in *Aedes aegypti*^68,69^. 40S ribosomal protein S15 (*AAEL011656*) is incorporated into translating ribosomes recruited by orthoflaviviral RNA^70,71^. Thioredoxin peroxidase (*AAEL013528*) buffers the oxidative stress that accompanies viral replication^72^. Ribulose-5-phosphate-3-epimerase (*AAEL010505*) acts in the pentose phosphate pathway, into which Zika and Japanese encephalitis viruses divert host glucose in infected mosquito and vertebrate cells^73,74^. The fifth gene, *AAEL012060* encodes a brain-enriched precursor of the PBAN/pyrokinin family, a conserved class of insect neuropeptides with roles in feeding regulation, gut motility, and pheromone biosynthesis^75–77^. In our neurotranscriptome dataset, *AAEL012060* is enriched in the brain relative to antennae, rostrum, proboscis, maxillary palps, legs, abdominal tip, and ovaries, and this enrichment is stable across sugar-fed and post-blood-meal states. Whether *AAEL012060* upregulation specifically in virus-positive heads, against a background of otherwise minimal infection-specific change, reflects a direct viral effect, a homeostatic response, or an unrelated change in feeding-state physiology cannot be resolved from these data, and behavioral consequences would require targeted functional studies.

None of the responses we observed correspond to canonical antiviral pathways, consistent with the absence of differential expression across 413 curated immunity genes (**Table S5, Data S8**), and the modest fold-changes could be a signature of low-level metabolic accommodation of viral replication rather than active defense.

The minimal transcriptional response of the brain contrasts with mosquito midguts and salivary glands, where arbovirus infection induces hundreds of differentially expressed genes including canonical immune effectors^13,16,36,43^. Several non-mutually-exclusive mechanisms could account for this difference. First, the *Aedes aegypti* brain is enclosed by a glial blood-brain barrier formed by perineurial and subperineurial glial cells, which in insects restrict access to hemolymph-borne pathogens, but also to immune effectors, and circulating hemocytes to the neural parenchyma^78,79^. This barrier may prevent the brain from sensing infection in the same way as the midgut and salivary glands. Second, neurons themselves may have a reduced capacity to mount canonical innate immune responses; in *Drosophila melanogaster*, immune signaling pathways are constitutively dampened in the central nervous system, in part because immune effectors tolerated in other tissues cause neuronal toxicity^80,81^. Third, the brain may rely on disease-tolerance mechanisms, physiological responses that limit damage from infection without reducing pathogen load, rather than on resistance mechanisms (active pathogen clearance) that dominate in the midgut^82^. Tolerance is energetically less costly and avoids immunopathology, both attractive properties for a tissue whose function depends on intact, non-renewable cells^82,83^. Fourth, DENV may actively suppress mosquito transcriptional responses through viral antagonists of mosquito immunity, as observed for orthoflaviviruses in other tissues and in cells^42,84,85^. Orthoflaviviral nonstructural proteins and subgenomic RNAs interfere with RNAi and other antiviral pathways and could account for the absence of detectable immune-gene upregulation. These mechanisms (host tolerance and neural protection, and virus-mediated suppression) are not mutually exclusive. Their combined effect may represent an evolutionary equilibrium in which both vector and virus benefit from neural tolerance: the mosquito avoids immunopathology in tissue critical for behavior and reproduction, and the virus preserves the host-seeking behavior on which its transmission depends.

The minimal transcriptional response of the mosquito brain also contrasts with vertebrate neural tissue. Although neuroinvasion is an uncommon complication of dengue in humans, it can produce encephalitis, neuronal damage, and death^44–46^. Experimental mouse models can show aspects of this neuropathology: RNA-sequencing of DENV-infected mouse brains shows neuronal apoptosis, immune cell infiltration, and MHC-I upregulation^86^, and bulk RNA-seq shows robust pro-inflammatory cytokine induction and brain damage^87^. These models rely on routes and host backgrounds that diverge from natural infection, including immunocompromised mice and direct viral inoculation, so the responses they elicit should be interpreted with caution and likely reflect both direct neural infection and the systemic immune dysregulation inherent to these systems. Where DENV does replicate in vertebrate neural tissue, it can drive immune activation, neuronal apoptosis, and cytokine induction. By contrast, productive DENV infection of the mosquito brain triggers no more than five differentially expressed genes, none corresponding to canonical immune pathways.

### Limitations of this study

We used a single DENV isolate belonging to serotype 1 and a single mosquito strain (Liverpool). Whether other virus isolates, serotypes, or vector genotypes show similar transcriptional dynamics remains to be tested. We sequenced whole heads with head appendages removed, so our data capture gene expression from multiple head structures, including the compound eyes, and cannot be attributed solely to the brain. We sampled three time points (7, 11, and 14 DPF) and may have missed transient transcriptional responses during the initial phase of infection. The number of virus-positive heads was limited (n=4 at 7 DPF, n=3 at 11 DPF under the strict head-only criterion), but concordance of PCA, unsupervised clustering, and differential expression analyses across grouping criteria nevertheless supports the conclusion that infection-specific transcriptional changes are minimal.

### Concluding remarks

The mosquito brain is permissive to DENV replication yet remains transcriptionally stable, a response distinct from that reported in midgut and salivary glands and from that of vertebrate neural tissue. Three questions follow directly from this work. What determines the bimodal infection pattern, and which cell types support viral replication in the brain? Does *AAEL012060* upregulation translate to changes in PBAN/pyrokinin signaling and to altered feeding or host-seeking behavior? And is brain transcriptional homeostasis a feature specific to DENV, or a general property of arboviral infection of mosquito neural tissue?

## METHODS

### Mosquito rearing and maintenance

The *Aedes aegypti* wild-type Liverpool (LVP) strain^48^ has been maintained in the Vosshall Laboratory insectary at The Rockefeller University since 2011 at 26-28°C and 70-80% relative humidity with a 14:10 h light:dark cycle. Approximately 3,000 LVP eggs were collected on wet Whatman qualitative paper (Sigma-Aldrich, WHA1001813) and shipped from Rockefeller University to the Institut Pasteur in Paris, where about 1,600 eggs were used to establish a laboratory colony.

Eggs were synchronously hatched by placing them in 100 mL of dechlorinated water (centrally filtered at the Institut Pasteur) in a vacuum chamber for 1 h. Larvae were reared in 24 × 34 × 9 cm plastic pans containing 1.5 L of dechlorinated water and fed daily with Tetramin fish food (Tetra). Larval density was maintained at 200 larvae per pan, with addition of water, until the pupal stage. Pupae were collected and transferred to 30 × 30 × 30 cm clear plastic insect rearing BugDorm cages (MegaView Science, DP1000). Upon emergence, adult mosquitoes were provided with sterile 10% sucrose solution (w/v in deionized water) *ad libitum*, contained in a 100 mL Pyrex Erlenmeyer flask and dispensed through a cotton wick. All larvae, pupae, and adults were maintained at 28°C, 70% relative humidity and a 12:12 h light:dark cycle. Mosquitoes in the Arthropod Containment Level 3 (ACL-3) laboratory were maintained in two incubators (Memmert, HPP260) set at the same temperature and humidity and light:dark cycle as the insectary.

The Institut Pasteur LVP colony was maintained by feeding adult female mosquitoes fresh commercial rabbit blood (BCL, Boisset St Priest) using a Hemotek membrane feeding apparatus (Hemotek Ltd, PS6220) with pig intestine mucosa (Tom Press, Sorèze) as the membrane. Mosquitoes were fasted for 18-20 h before blood feeding and allowed to feed for approximately 30 min. Humid blotting paper for egg laying (Whatman, WHA1001813) was placed in each cage one day post-blood feeding, and eggs were collected 10-14 DPF. Collected eggs were dried and stored at room temperature before hatching to establish the next generation.

### DENV production

Mosquitoes were experimentally infected with the dengue virus serotype 1 (DENV-1) isolate KDH0026A (GenBank: HG316481.1), originally isolated in 2010 from the serum of an infected patient from Kamphaeng Phet, Thailand^49^. Virus stocks were produced in *Aedes albopictus*-derived C6/36 cells (Sigma-Aldrich, 89051705) grown in Leibovitz’s L-15 medium (Thermo Fisher Scientific, 11415064) supplemented with 10% fetal bovine serum (FBS; Thermo Fisher Scientific, A5256701), 1× non-essential amino acids (Thermo Fisher Scientific, 11140050), and 0.1% penicillin-streptomycin (Thermo Fisher Scientific, 15140148) at 28°C. Cell culture supernatants were harvested, supplemented with 10% FBS and sodium bicarbonate (pH ∼8), aliquoted, and stored at −80°C. The same virus stock was used for all infections described in this paper. Virus concentration was quantified by focus-forming assay (FFA) as described below.

### Focus-forming assay

DENV titers were determined by FFA as previously described^88^. C6/36 mosquito cells^89^ were seeded at a density of 5 × 10⁴ cells/well in 96-well plates 24 h prior to inoculation. Serial dilutions of virus samples were prepared in Leibovitz’s L-15 medium supplemented with 0.1% penicillin-streptomycin, 2% tryptose phosphate broth (Thermo Fisher Scientific, 18050039), 1× non-essential amino acids, and 2% FBS. Cells were inoculated with 40 μL of diluted sample and incubated for 1 h at 28°C. Following incubation, the inoculum was replaced with 150 μL of overlay medium consisting of a 1:1 mixture of supplemented Leibovitz’s L-15 medium and 1.6% carboxymethylcellulose (Sigma-Aldrich, C4888), with 2× Antibiotic-Antimycotic (Thermo Fisher Scientific, 15240062) and 10% FBS.

Plates were incubated at 28°C for 5 days. Cells were fixed for 30 min with 3.6% paraformaldehyde (Sigma-Aldrich, 158127), washed three times with 1× phosphate-buffered saline (PBS; Thermo Fisher Scientific, 10010031), and permeabilized for 30 min with 0.3% Triton X-100 (Sigma-Aldrich, X100) in PBS at room temperature. After three PBS washes, cells were incubated for 1 h at 37°C with mouse anti-DENV complex monoclonal antibody (Sigma-Aldrich, MAB8705) diluted 1:200 in PBS containing 1% bovine serum albumin (BSA; Interchim, UPQ84170). Following three additional PBS washes, cells were incubated for 30 min at 37°C with Alexa Fluor 488-conjugated goat anti-mouse antibody (Thermo Fisher Scientific, A11001) diluted 1:500 in PBS with 1% BSA. After final washes in PBS and water, infectious foci were visualized and counted using an Evos FL digital fluorescence microscope (Thermo Fisher Scientific).

### Mosquito oral exposure to DENV

Mosquitoes were orally exposed to DENV following established feeding protocols^88^. *Aedes aegypti* LVP mosquitoes (5-7 days old) were cold-anesthetized for sex separation, and groups of 50 females were transferred in 1-pint cardboard soup boxes, moved to an incubator within the ACL-3 laboratory and deprived of sugar for 24 h prior to experimental infection.

Artificial meals were prepared fresh the morning of each experimental day. Fresh rabbit whole blood (BCL, Boisset St Priest) was centrifuged for 15 min at 1,200 rpm in an Eppendorf 5920 Centrifuge (Sigma-Aldrich, EP5948000010) to separate erythrocytes from plasma. Erythrocytes were washed three times by resuspending them in 1× PBS (Thermo Fisher Scientific, 10010031) and centrifuging for 5 min at 1,400 rpm. Following final centrifugation, the basic meal was prepared by resuspending rabbit erythrocytes to the original blood volume in 1× PBS supplemented with 10 mM adenosine triphosphate (Sigma-Aldrich, A2383) as a phagostimulant.

Mosquitoes were fed one of the following three meals: (1) 2:1 volumetric ratio of basic meal to DENV stock with a viral titer of 10⁶ FFU/mL; or (2) 2:1 volumetric ratio of basic meal to heat-inactivated DENV prepared by heating at 56°C for 30 min in an Eppendorf Mastercycler Nexus Gradient Thermocycler (Thermo Fisher Scientific, 12304943) to inactivate the virus without denaturing the viral proteins^50^; or (3) 2:1 volumetric ratio of basic meal to supernatant from uninfected C6/36 cells. Aliquots of all three meals were tested in the FFA to confirm the presence or absence of DENV.

Groups of 50 female mosquitoes were allowed to feed for 20 min through an artificial membrane-feeding system as previously described. After feeding, mosquitoes were anesthetized at 4°C for 10-15 min and sorted on ice to isolate fully engorged females by visual inspection. Groups of 30-40 fed-females were maintained in 1-pint cardboard soup boxes under standard insectary conditions with permanent access to 10% sucrose solution and moist oviposition paper (Whatman, WHA1001813). All mosquitoes, regardless of meal type, were maintained in the ACL-3 laboratory.

### Tissue collection

Mosquitoes were anesthetized at 4°C for 15-20 min and dissected in ice-cold RNase-free 1× PBS (Thermo Fisher Scientific, 10010031) under a stereomicroscope (Leica, M60) in the ACL-3 laboratory. Heads were severed using either a 0.15 mm micro knife (Fine Science Tools, 10316-14) or a straight 0.175 mm micro scissors (Fine Science Tools, 15030-14). Head appendages (antennae, maxillary palps, proboscis), carcasses, and legs were removed with Dumont #5 forceps (Fine Science Tools, 11251-20). All tools were cleaned with 70% ethanol and RNase Away (Thermo Fisher Scientific 7002PK) between dissections, using a dedicated set of tools for each experimental condition.

Heads designated for whole-brain immunostaining were cut and transferred individually to 24-well cell culture plates (Sigma-Aldrich, CLS353226) and incubated with rotation in a fixative solution containing 0.25% Triton X-100 (Sigma-Aldrich, X100) and 4% paraformaldehyde in 0.1 M phosphate-buffered saline (pH 7.4; Electron Microscopy Sciences, 15735-85) at 4°C for 3 h. Following fixation, heads were washed three times in sterile 1× PBS and stored at 4°C. Plates containing fixed mosquito heads in 1× PBS were shipped from Institut Pasteur to Rockefeller University at 4°C using a specialized courier service with all appropriate import permits in place (World Courier France) for brain dissection and imaging.

Heads designated for individual bulk RNA sequencing, and carcasses and pools of legs from all the mosquitoes we dissected, were transferred to vials containing sterile, RNase-free glass beads and ice-cold 250 μL TRIzol (Thermo Fisher Scientific, 15596026). Heads designated for the preliminary pooled bulk RNA-seq were placed individually in 96-well plates (Sigma-Aldrich, CLS3367), then pooled in groups of 10 per tube following the RT-qPCR validation described below. All collected tissues were kept on wet ice, dissected in batches of ∼10, then homogenized with a Precellys instrument (Bertin, P000669-PR240-A.0) for 60 s and immediately stored in a −80°C freezer.

### Whole brain staining and imaging

Mosquito brains were dissected and immunostained at Rockefeller University in New York, USA, as described previously^90^. Briefly, brains were dissected from paraformaldehyde-fixed mosquito heads using sharpened Dumont #5 forceps (Fine Science Tools, 11251-20) and placed individually in Falcon cell-strainer caps (Sigma-Aldrich, CLS431751) within 24-well plates (Sigma-Aldrich, CLS353226) filled with 1× PBS (Thermo Fisher Scientific, 10010031). Brains underwent six 15-min washes in PBT (0.25% Triton X-100 [Sigma-Aldrich, X100] in 1× Phosphate-Buffered Saline) at room temperature, followed by permeabilization in 4% Triton X-100 with 2% normal goat serum (Thermo Fisher Scientific, 31872) in 1× PBS for 48 hours at 4°C. After six additional 15-min washes in 0.25% PBT, brains were incubated for 48 hours at 4°C with the primary antibodies: mouse anti-DENV complex (Sigma-Aldrich, MAB8705) and rat anti-N-Cadherin (DSHB, ADL67.10), both diluted 1:100 in 0.25% PBT with 2% normal goat serum. Following six 15-min washes in 0.25% PBT, brains were incubated with NucBlue Fixed Cell ReadyProbes Reagent (Thermo Fisher Scientific, R37606) and secondary antibodies: Goat anti-Rat Alexa Fluor Plus 488 (Thermo Fisher Scientific, A48262) and goat anti-Mouse Alexa Fluor Plus 555 (Thermo Fisher Scientific, A32727) diluted 1:200 in 0.25% PBT with 2% normal goat serum for 48 h at 4°C. After six final 15-min washes in 0.25% PBT, brains were mounted in SlowFade Diamond (Thermo Fisher Scientific, S36967) using #1.5 coverslips (Sigma-Aldrich, CLS2975245) as spacers for confocal imaging. All the staining and washing steps were performed on a low-speed orbital shaker.

Images were acquired using an Inverted Zeiss Axio Observer Z1 LSM 880 NLO laser scanning confocal microscope and processed with the Zeiss ZEN software (V. 3.10). A Plan-Apochromat 25×/0.8 glycerol immersion-corrected objective (Zeiss, 420852-9870-790) at 1024×1024 pixel resolution was used to capture whole-brain images with 4× averaging. We optimized confocal settings for qualitative detection of fluorophores, precluding quantitative intensity analysis, and image acquisition settings were kept consistent across all the experiments. When necessary, tiled images were stitched with 10% overlap. High-magnification images of specific brain regions were captured with a Plan-Apochromat 63×/1.4 oil objective (Zeiss, 420782-9900-799).

Brains were scored by visual inspection of maximum-intensity projection images. “No cells” brains contained no detectable DENV antigen signal above background. “Few cells” brains contained fewer than 15 isolated DENV-positive cells per brain, located at the brain periphery. “Many cells” brains showed multiple, usually more than 100, DENV-positive cells distributed across multiple brain regions. No brains fell between these categories, consistent with the bimodal infection pattern described in the Results. All scoring was performed by a single observer (U.P.).

### RT-qPCR on carcasses and legs

Total RNA from legs and carcasses was extracted using the NucleoSpin 96 RNA Extraction Kit (Macherey-Nagel, 740709.4) according to manufacturer instructions, with TRIzol (Thermo Fisher Scientific, 15596026) substituted for the provided lysis buffer. Carcasses and pools of legs were eluted in 50 μL and 30 μL of RT-PCR grade DNase/RNase-free Water (Thermo Fisher Scientific, AM9935), respectively. Extracted RNA samples and DENV RNA standards were shipped to Rockefeller

University at −80°C using a medical courier with all appropriate import permits in place (Marken France). cDNA was synthesized using the GoScript Reverse Transcription System (Promega, A2801). Multiplex qPCR was performed using the GoTaq Probe qPCR System (Promega, A6101). We used probes and primers targeting the DENV *NS5* gene and two *Aedes aegypti* housekeeping genes tested in the literature^53^: *Actin5c* (*AAEL011197*) and *RPS17* (*AAEL004175*). Primers and probe concentrations were 250 nM and 75 nM respectively. Each qPCR experiment included triplicate biological replicates of each sample, either cDNA from DENV-exposed mosquito legs or carcasses and two negative controls: no-reverse transcriptase and no-template controls. For absolute quantification of viral copies, a 5-point standard curve was generated using cDNA synthesized with the GoScript Reverse Transcription System from 10-fold serial dilutions ranging from 10^7^ to 10^2^ DENV copies. qPCR was performed using an Applied Biosystem QuantStudio 5 instrument (Thermo Fisher Scientific, A34322). Raw data were analyzed with the QuantStudio Design & Analysis Software v 1.6.1. Viral reads were normalized to the expression of two housekeeping genes with the 2^ΔΔCq^ method implemented in the NormqPCR package^91^ version 1.48.0, running on R version 4.3.3.

Mosquito legs and carcasses were classified as positive for infection when RT-qPCR exceeded 10 DENV genome copies/µL. This RT-qPCR–based threshold was used only to calculate population infection rates and is distinct from the read-count criterion used to assign RNA-seq group membership (>100 DENV reads per library; see above). Infection rates at each time point are reported with 95% Wilson score confidence intervals. Differences in infection rates across time points were assessed by chi-square test, with pairwise comparisons by Fisher’s exact test. All data are available in **Table S2**.

### Total RNA library preparation and sequencing

Total RNA was extracted from individual or pooled *Aedes aegypti* female heads (without head appendages) at the Institut Pasteur using the PicoPure Kit (Thermo Fisher Scientific, KIT0204) with the following modifications: TRIzol lysates were thawed and incubated at room temperature for 5 min and then transferred to pre-centrifuged PhaseLock tubes (Thermo Fisher Scientific, A33248) containing 48 µL of chloroform:isoamyl alcohol 24:1 (Sigma-Aldrich, C0549-1PT). The mixture was shaken by hand for 30 s, incubated for 2 min, and centrifuged at 12,000 rpm for 15 min at 4°C in a Beckman Coulter Allegra X-12 benchtop refrigerated centrifuge. The aqueous TRIzol layer was then pipetted onto the PicoPure silica column, and subsequent steps were performed according to the manufacturer’s PicoPure protocol, including treatment with RNase-free DNase (Qiagen, 79254). RNA samples were stored at −80°C at Institut Pasteur and shipped to Rockefeller University on dry ice using a specialized medical courier service with all appropriate import permits in place (Marken France) with guaranteed monitoring and dry ice replenishment.

RNA samples were assessed for quality and concentration using a Bioanalyzer 2100 instrument (Agilent, G2939B) with the RNA Pico kit (Agilent, 5067-1513). Samples with RNA concentrations below 1 ng/µL were concentrated using a Vacufuge (Eppendorf, EP5305000169) for 10 min at 30°C. RNA libraries from individual heads were prepared by integrating the Zymo-Seq RiboFree Total RNA Library Kit (Zymo Research, R3003) for ribosomal RNA depletion and the TruSeq Stranded Total RNA Library Prep Kit (Illumina, 20020597) for cDNA synthesis, fragmentation, and adapter ligation. RNA libraries from pooled mosquito heads were prepared with the TruSeq Stranded mRNA library prep kit (Illumina, 20020595) and so underwent poly(A) enrichment instead of ribosomal RNA depletion. The RNA libraries were sequenced on an Illumina NovaSeq 6000 instrument using an S2 flowcell (Illumina, 20028314) for the rRNA-depleted libraries and an S1 flowcell (Illumina, 20028317) for the poly(A)-enriched libraries. Libraries were sequenced to a depth of 30-40 million paired 100-bp short reads per sample. Raw reads are available on SRA under the BioProject ID PRJNA1321547.

### RNA-sequencing data quality control and mapping

All bioinformatics analysis was performed with computational resources provided by the Rockefeller University High Performance Computing Resource Center (RRID:SCR_025889). We ran the nextflow-based nf-core/rnaseq^92,93^ pipeline (version 3.17.0) in a dedicated Conda environment, configured with Nextflow^94^ v24.10.9, to quantify mosquito gene expression. The pipeline performed quality control using FastQC (v0.12.1). Subsequently, adapter sequences were removed, and reads were quality-trimmed using Trim Galore! (v0.6.10). Genomic contaminants and ribosomal RNA remnants were removed using SortMeRNA (v4.3.7). Reads were mapped to the AaegL5.0 *Aedes aegypti* reference genome^65^ (AaegyptiLVP_AGWG, downloaded from VectorBase release 68) using STAR (v2.7.11b) with mapping rates of 75-90%. Downstream alignment quantification was performed using Salmon (version 1.10.3). Duplicate reads were marked using Picard MarkDuplicates (v3.1.1). Transcript assembly and quantification were performed using StringTie (v2.2.3), and genomic coverage was visualized through bigWig files using BEDTools (v2.31.1) and bedGraphToBigWig. Comprehensive quality assessment was conducted using multiple tools: RSeQC (v5.0.2) and Qualimap (v2.3) for general sequencing metrics, dupRadar (v.1.32.0) for duplication rate analysis and DESeq2 (v1.34) for expression-level diagnostics. Unaligned sequences were taxonomically classified using Kraken2 (v2.1.3) with abundance estimation refined by Bracken (v2.9). Kraken2 was run against the entire RefSeq viral database (https://www.ncbi.nlm.nih.gov/labs/virus; September 4, 2024, release) to detect DENV reads with precision. To validate Kraken2-Bracken results, the nf-core/viralrecon^93,95^ pipeline (version 2.3) was used to map RNA-seq reads to our DENV DV26A isolate genome and perform de novo viral genome assembly using bowtie2 (v2.4.4) and Mosdepth (v0.3.3) to calculate viral genomes coverage.

### Differential Gene Expression analysis

Raw, length-scaled counts from the nfcore/rnaseq pipeline were initially processed using the nf-core/differentialabundance^93,96^ pipeline (version 1.5.0; **Data S5 and S6**). This pipeline cross-checks input matrices, sample annotations, and contrasts for consistency, followed by differential expression analysis across all specified contrasts using DESeq2^64^ (v1.34). Quality assessment and exploratory analysis were performed using PCA, sample clustering, and expression visualization tools.

To address potential outliers, a common issue with RNA-seq data, while maintaining compatibility with DESeq2, we initially tried a percentile-based winsorization approach with a custom R script based on an existing method^97^. Raw counts were first normalized using DESeq2^64^ (v1.34) size factors, extreme values above the 93rd percentile were replaced with the 93rd percentile value for each gene, and the data were converted back to integer counts by multiplying by size factors and rounding. This approach removed false positives but also true positives because replicates were not enough to efficiently calculate percentiles, and removing variation could hurt statistical power.

### Post-DESeq2 outlier and artifact annotation of differentially expressed genes

To minimize false-positive differentially expressed genes, we applied a post-hoc annotation to the significant DEGs (DESeq2, padj < 0.01) that labeled each gene Clean, Outliers_Present, or Artifact based on whether its statistical significance was supported by a group-level pattern or by single-sample extremes (Github: https://github.com/UmbertoPalatini/DeSeq2_DEGs_OutlierDetector). Two rules were applied in sequence. Rule 1 (single-sample outlier detection) examined each gene’s normalized-count vector under four complementary criteria: ratio to the sample median (>20×), sparsity-aware ratio to the sample mean for zero-median vectors (>10×), ratio to the smallest non-zero value (>1000×), and a robust 3×IQR criterion. A gene was classified as Artifact when ≤2 outliers were detected, ≥2 non-outlier samples remained per group, all outliers fell within a single contrast group, and at least one magnitude threshold was met (normalized counts >10,000; >100× group median; or >50× median combined with >10× the maximum non-outlier value and >1,000 counts). Genes with outliers but a coherent group-level pattern were retained as Outliers_Present. Rule 2 (sparse expression) flagged any DEG with non-zero counts in fewer than three samples across the contrast as Artifact; complete-induction patterns (zero in one group, expressed in all replicates of the other) exceeded this threshold and were retained as Clean. The procedure was calibrated to flag extreme technical artifacts (single sample expression spikes ∼100-1000x above background) while preserving biologically relevant fold-changes (2–20x), and complements DESeq2’s Cook’s-distance filtering, which does not act when the smaller contrast group has ≤3 replicates. Artifact-flagged and Outliers_Present genes were excluded from downstream analyses; only Clean genes were retained. The full list of DEGs, including artifact-flagged and Outliers_Present genes, is available in the supplementary data files.

### Data visualization and statistics

Pairwise comparisons between experimental groups (DENV-exposed vs no virus, DENV-exposed vs heat-inactivated, heat-inactivated vs no virus) to determine differential expression significance were conducted using the Wald test implemented in DESeq2, with adjusted p-values calculated using the Benjamini-Hochberg procedure to control the false discovery rate. Gene expression heatmaps were generated by calculating row-wise Z-score normalization of DESeq2-normalized counts, which centers expression values around zero and scales by standard deviation to enable visualization of relative expression patterns across samples. Hierarchical clustering of genes and samples was performed using Ward’s minimum variance method based on Euclidean distances. Heatmaps were generated using the pheatmap package^98^ (version 1.0.13) in R (version 4.2.1). Principal component analyses (PCA) were performed on variance-stabilized transformed counts using the plotPCA function from DESeq2^64^ to assess global transcriptional patterns and identify major sources of variation across experimental groups and time points. Data visualization was performed using a combination of R and Python packages. Expression bar plots, volcano plots, and dot plots were generated using ggplot2^99^ (version 3.4.0) within the tidyverse framework (version 2.0.0) in R, with statistical summaries calculated using the stats package (version 4.2.1). Additional visualizations and exploratory analyses were conducted in Python using matplotlib^100^ (version 3.10.0), seaborn^101^ (version 0.13.1), pandas^102^ (version 2.3.0), and numpy^103^ (version 2.3.0). Figures were created in Adobe Illustrator.

## DATA AVAILABILITY

Raw data produced in this study are available at NCBI SRA, EBI-ENA and DDBJ databases under BioProject PRJNA1321547. Raw bulk RNA-seq data from the mosquito neurotranscriptome atlas previously published and re-analyzed in this study are under BioProject PRJNA236239. Additional raw data, including normalized RNA expression counts, are available in Data File S1. The custom script for identifying outliers is available on Github: https://github.com/UmbertoPalatini/DeSeq2_DEGs_OutlierDetector.

## ACKNOWLEDGEMENTS

We thank members of the Vosshall Lab at The Rockefeller University for comments on the manuscript; Laura Duvall for insights into PBAN neuropeptide biology; Margaret Herre for advice and technical guidance on brain dissections and immunofluorescence staining; Libby Mejia and Melissa Dallesandro for expert mosquito rearing in the Vosshall Lab; Paula Calle for laboratory management and logistic support. We also thank all members of the Lambrechts lab at Institut Pasteur for their help and support; Catherine Lallemand for mosquito rearing in the Lambrechts Lab; Elodie Couderc, Alicia Lecuyer, Thomas Vial, and Shiho Torii for support with experimental protocols and Théo Maire for fruitful discussion and fostering of ideas. We thank the following resource centers at Rockefeller: Connie Zhao and Bin Zhang at the Rockefeller Genomics Resource Center (RRID:SCR_020986) for setting up a new protocol for rRNA depletion of total RNA; Alison North, Priyam Banerjee, and Christina Pyrgaki at the Rockefeller Bio-Imaging Resource Center (RRID:SCR_017791) for their support and assistance with confocal imaging.

## AUTHOR CONTRIBUTIONS

**U.P.** conceived the study, designed and performed all infections and dissections at the Institut Pasteur in Paris with the help of **S.D.**, performed all immunofluorescent staining and imaging, produced and analyzed the RNA-seq data and wrote the manuscript. **S.D.** prepared the DENV stock, performed the viral titrations, and assisted with mosquito rearing and DENV infections at the Institut Pasteur. **A.R.-V.** and **Y.N.T.** carried out all mosquito brain dissections and mounted the immunostained samples for confocal microscopy. **A.E.D.** produced figures and supported manuscript writing. **N.S.** supported manuscript writing and managed experiments. **L.L.** supervised the study and designed ACL-3 experiments, provided viral stocks and resources, and secured funding. **L.B.V.** conceived the study alongside U.P., supervised the study, secured funding, and wrote the manuscript with **U.P.**

## FUNDING

This research was supported by the Stavros Niarchos Foundation (SNF) as part of its grant to the SNF Institute for Global Infectious Disease Research at The Rockefeller University, an EMBO Long-Term Fellowship (ALTF 664-2022) and a Human Frontier Science Program fellowship (LT 0012-2023L) (U.P.). N.S. was supported by an EMBO Long-Term Fellowship (ALTF 286-2019). L.B.V. is supported by the Howard Hughes Medical Institute. L.L. was supported by the French Government’s Investissement d’Avenir program through the Laboratoire d’Excellence Integrative Biology of Emerging Infectious Diseases (grant ANR-10-LABX-62-IBEID) and the Inception program (grant ANR-16-CONV-0005).

## CONFLICTS OF INTEREST

The authors declare no competing interests.

## SUPPLEMENTARY FIGURES

**Supplementary Figure 1.**
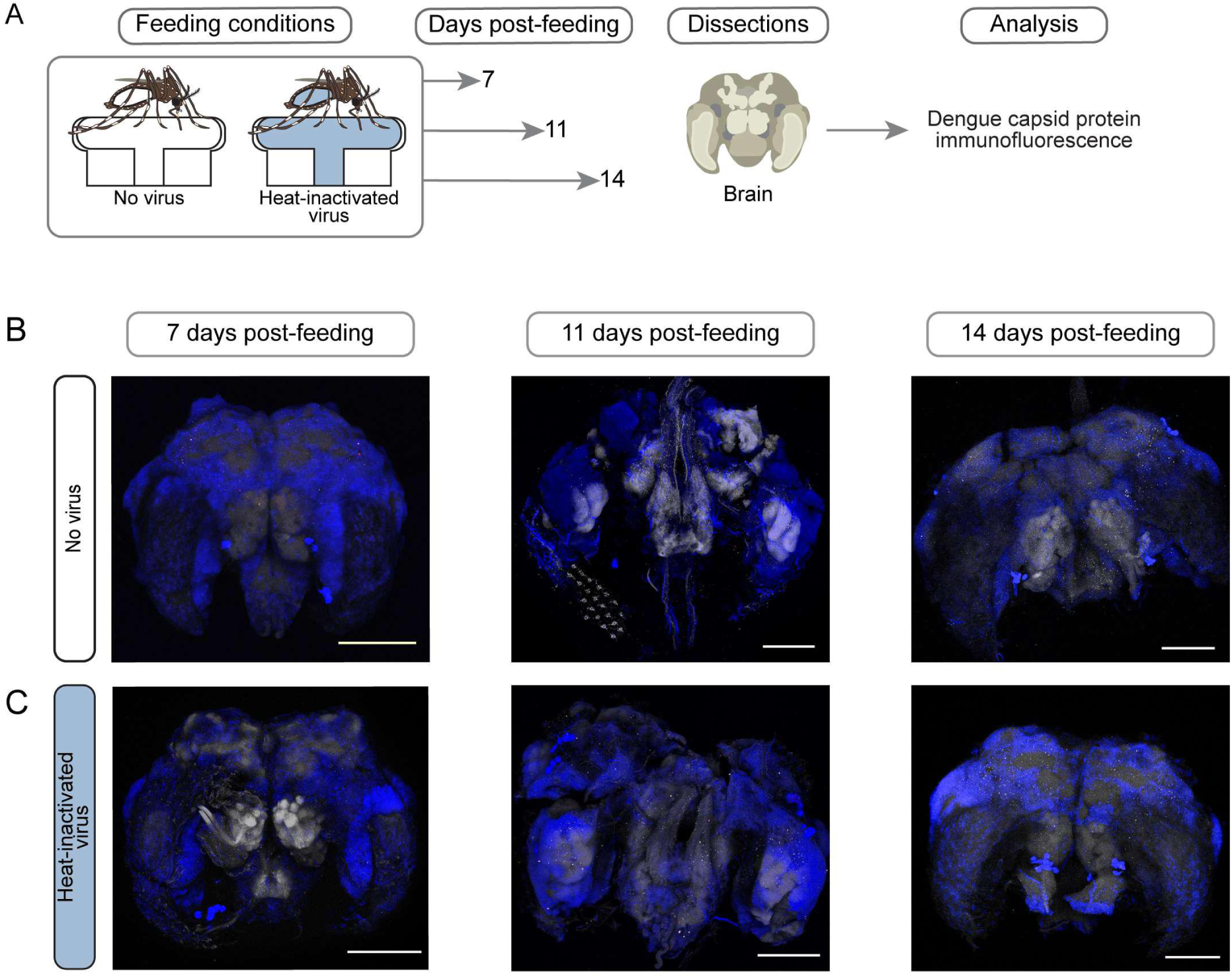
| Brains of mosquitoes fed heat-inactivated virus show no DENV complex antigen immunoreactivity. (**A**) Experimental workflow. (**B, C**) Representative maximum-intensity projections of whole-mount brains from mosquitoes fed no virus (**B**) or heat-inactivated virus (**C**) and stained for DENV antigen using an anti-DENV complex antibody (magenta), anti-N-cadherin (grey), and DAPI (blue) (n = 16 per condition). Scale bars, 100 μm. See also **Table S2**.

**Supplementary Figure 2.**
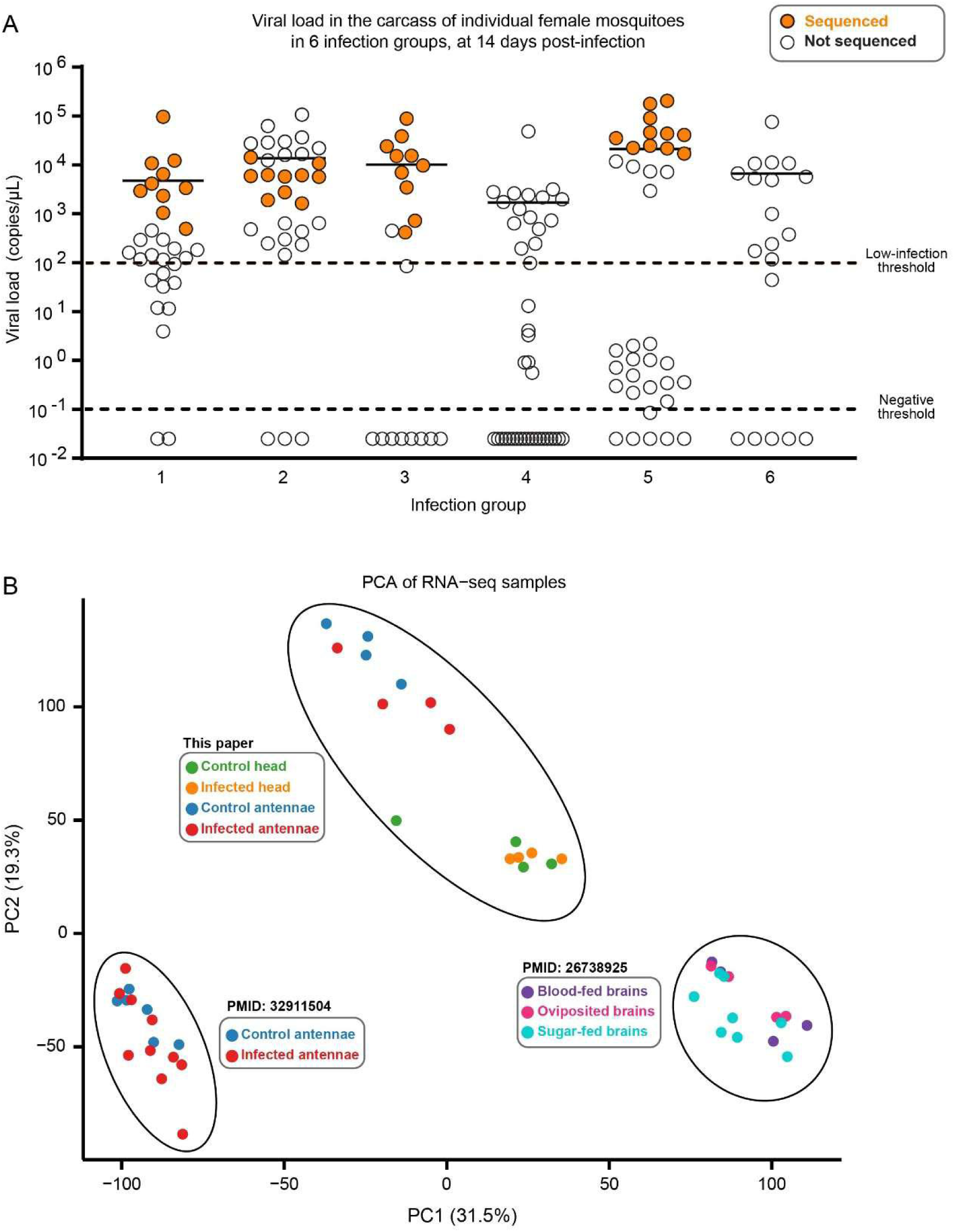
| Preliminary pooled RNA-seq experiment. **(A)** DENV RNA copies per carcass in individual mosquitoes at 14 DPF, measured by RT-qPCR. Each circle represents one mosquito across six independent infection groups. Horizontal black bars indicate group medians. Dashed lines mark the low-infection threshold (10^2^ copies/μL) and negative threshold (10^-1^ copies/μL). Filled orange circles indicate heads selected for pooled RNA-seq within an infection group, and open circles indicate individuals that were not sequenced. See also **Table S3**. **(B)** Principal Component Analysis (PCA) of RNA-seq data comparing pooled head and antennae samples to published *Aedes aegypti* brain and antennae datasets^33,67^. Data from each source are contained in an ellipse for clarity.

**Supplementary Figure 3.**
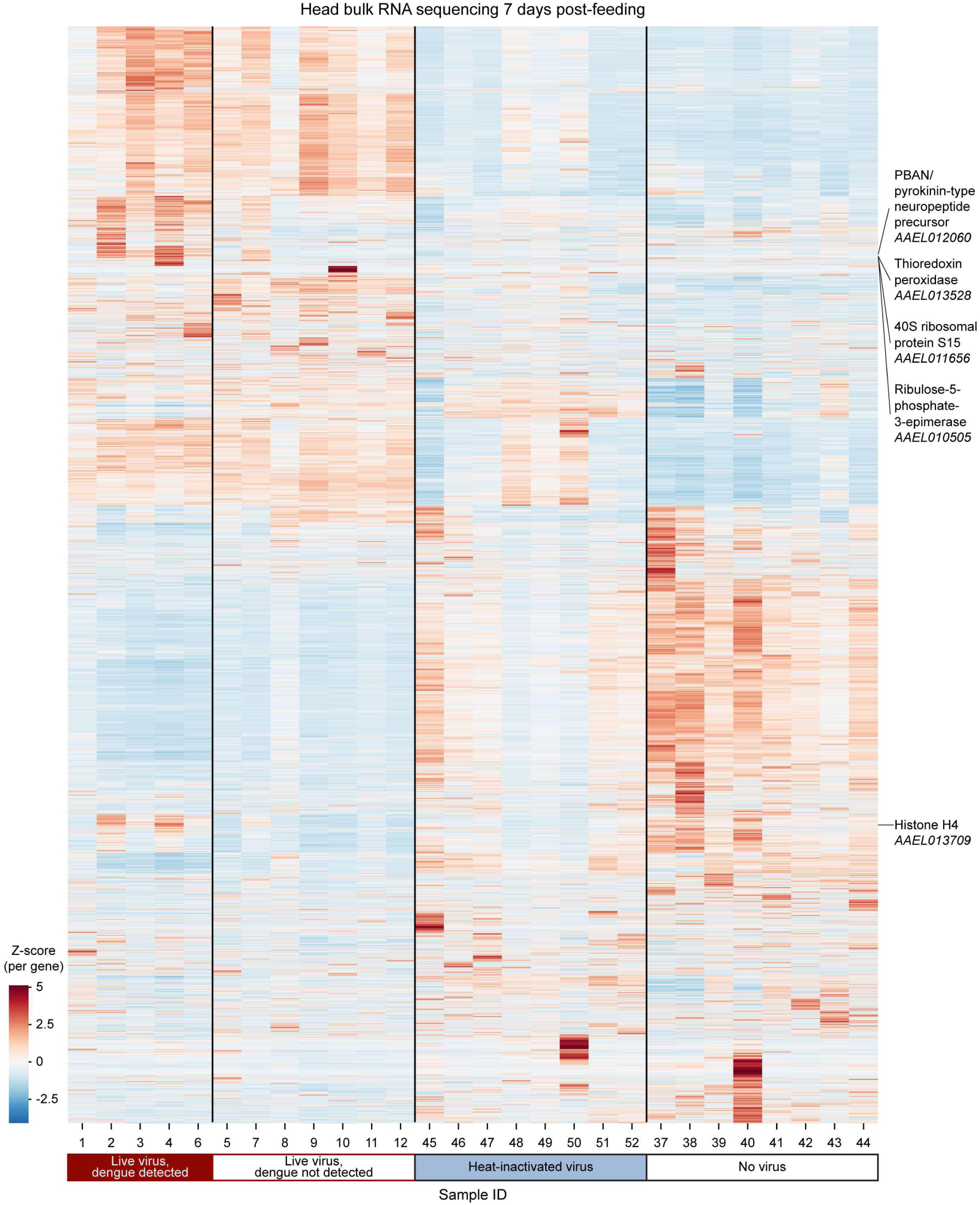
| Heatmap of gene expression in individual mosquito heads at 7 DPF. Each column represents one sequenced mosquito head; rows represent the 13,272 genes expressed in mosquito heads hierarchically clustered using Ward’s method (**Data S5**). Samples are classified as virus-positive or virus-negative based on the more comprehensive criterion (>100 viral reads in head and/or carcass). Colors indicate per-gene Z-scores of normalized expression values. Vertical lines separate experimental conditions. The five differentially expressed genes identified between virus-positive or virus-negative heads (padj < 0.01, |log₂FC| ≥ 1; see Figure 4) are labeled; *AAEL021837* is excluded because its significance derives from a mapping artifact rather than a biologically meaningful change

**Supplementary Figure 4.**
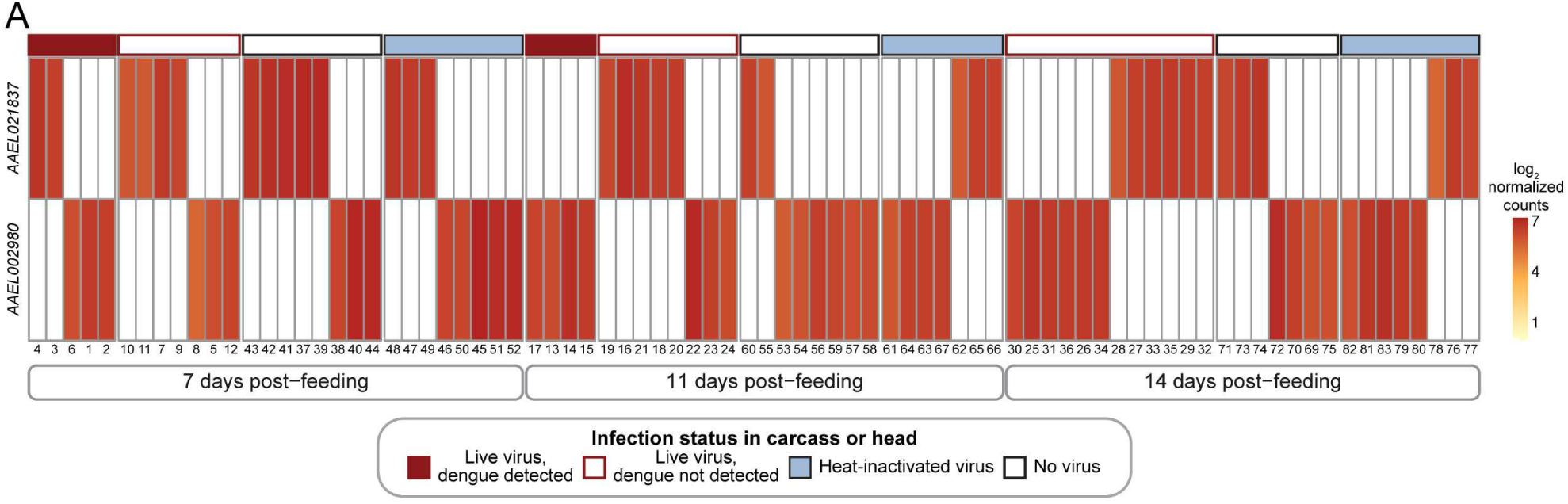
| Paralog read-assignment artifact at the *AAEL021837*/*AAEL002980* locus. Heatmap of log₂-normalized expression counts for *AAEL021837* (top) and its paralog *AAEL002980* (bottom) across all 83 sequenced mosquito heads (**Data S5**). The two genes share 100% nucleotide identity over 930 bp and map to the same UniProt entry (Q16H40). Expression is mutually exclusive across samples (Pearson r = −0.90): reads assign entirely to one locus or the other depending on the sample, consistent with stochastic mapping of ambiguous reads between identical paralogs. The color bar indicates experimental condition. Samples IDs are indicated as in **Table S2**.

## SUPPLEMENTARY TABLES AND SUPPLEMENTARY DATA

**Table S1. | Comprehensive sample tracking and experimental allocation.** Overview of the full cohort of mosquitoes used in this study, classified by infection condition (live DENV, heat-inactivated DENV, no virus) and collection time point (7, 11, and 14 DPF). For each group, the table reports the number of animals prepared, successfully blood-fed, incubated, collected, stored as backup at Institut Pasteur, and shipped to Rockefeller University for downstream analysis. For DENV-exposed cohorts, individual viral loads were quantified by RT-qPCR. Samples shipped to Rockefeller University are further broken down by experimental allocation: RNA-seq (sequenced), RNA (extracted but not sequenced), immunofluorescence (IF) imaged, and IF not imaged.

**Table S2. | Individual sample identifiers and experimental metadata.** Per-mosquito metadata for all 419 animals tracked in this study. Each row corresponds to one individual, identified by a unique Mosquito_ID. Columns report the Library_ID and RNAseq_ID (where applicable), infection date, experimental condition (Live DENV, Heat-Inactivated DENV, No virus), collection date, days post feeding (DPF), experimental group (Main Experiment or Backup in Pasteur), and downstream experimental allocation (RNA sequencing, IF Imaging, Full Body). For DENV-exposed samples processed by RT-qPCR, DENV copy numbers per µl are reported separately for carcass and legs. Library_ID encodes the sequential library number, time point, and condition suffix (DV = live DENV, H = heat-inactivated DENV, N = no virus).

**Table S3. | DENV viral loads in individual mosquitoes used in the preliminary pooled RNA-seq experiment.** DENV RNA copies per microliter of cDNA quantified by RT-qPCR in the carcass of individual *Aedes aegypti* females at 14 DPF, from the preliminary pooled RNA-seq experiment (2023). Each column corresponds to one of six independent infection groups (separate blood-feeding experiments performed on different dates), and each cell reports the viral load of one individual mosquito within that group. Values are absolute DENV genome copies per microliter, derived from RT-qPCR Cq values normalized against *Actin-5c* (AAEL011197) and *RPS17* (AAEL004175) using a standard curve of in vitro–transcribed DENV-1 RNA. The value 0.025 corresponds to the assay’s lower limit of quantification (below which samples were assigned this floor value). Empty cells indicate the infection group had fewer individuals than the column above (groups had unequal sample sizes). These data refer to Supplementary Figure 2A.

**Table S4. | Differentially expressed gene counts per comparison.** Summary of differentially expressed genes (as per DESeq2, Wald test, Benjamini-Hochberg FDR) for every pairwise comparison performed on the individual-mosquito RNA-seq dataset. Counts are given both for the raw DESeq2 output and after the post-hoc outlier filter (sparse-expression rule; see Methods), and the filtered set is split into down- and up-regulated genes. Results are reported under two definitions of infection status: infection assigned from DENV reads in the head library, and infection assigned from DENV reads in the paired carcass library.

**Table S5. | Immunity gene differential-expression summary by category.** Counts of differentially expressed genes (DESeq2, Wald test, Benjamini-Hochberg FDR; padj < 0.01, |log₂FC| ≥ 1, after the post-hoc outlier filter) within a curated set of 413 immunity-relevant *Aedes aegypti* genes grouped into 27 functional categories, with infection status assigned from DENV reads in the head library. Gene-level calls for the full set of 461 immunity and brain genes are provided in Data S8.

**Data S1. | Normalized RNA-seq read counts, pooled heads.** Data from the preliminary pooled sample experiment. DESeq2 size-factor-normalized read counts for 14,957 genes across 8 pooled head RNA-seq libraries (poly(A)-enriched mRNA, 14 DPF): 4 pools of DENV-exposed heads (INF_head_*) and 4 pools of control heads (CTRL_head_*). The first column lists *Aedes aegypti* gene identifiers (AaegL5 / VectorBase release 68 annotation, including manually curated chemoreceptor genes); each remaining column is one pooled library. Values are normalized counts (not log-transformed).

**Data S2. | Normalized RNA-seq read counts, pooled antennae.** As Data S1 but for the RNA-seq data from the antennae. DESeq2 size-factor-normalized read counts for 16,587 genes across 8 pooled antennae RNA-seq libraries (14 DPF): 4 pools of DENV-exposed antennae (INF_ant_1–INF_ant_4) and 4 pools of control antennae (CTRL_ant_1–CTRL_ant_4). Row and value conventions match Data S1.

**Data S3. | Differentially expressed genes, pooled heads.** DEGs from the pooled head experiment: DENV-exposed vs control pools, 14 DPF (DESeq2, Wald test, Benjamini-Hochberg FDR; padj < 0.01, |log₂FC| ≥ 1). One row per gene, with annotation, the full DESeq2 statistics, and the post-hoc outlier-filter output (whether the gene is a clean DEG, carries outlier pools, or is flagged as a likely artifact).

**Data S4. | Differentially expressed genes, pooled antennae.** As Data S3, for the pooled antennae experiment (DENV-exposed vs control antennae pools, 14 DPF). One row per gene; column definitions are identical to Data S3.

**Data S5. | Normalized RNA-seq read counts, individual heads.** DESeq2 size-factor-normalized read counts for 13,272 genes across 83 individual head RNA-seq libraries (rRNA-depleted total RNA; all conditions and time points). The first column lists *Aedes aegypti* gene identifiers (AaegL5 / VectorBase genome annotation release 68, including the manually curated chemoreceptor genes); each remaining column is one head library, identified by its RNAseq_ID (the numeric sample index used in Table_S1_samples.xlsx). Values are normalized counts (not log-transformed); 0 indicates no reads assigned to that gene in that library.

**Data S6. | Normalized RNA-seq read counts, individual carcasses.** DESeq2 size-factor-normalized read counts for 14,899 genes across 36 individual carcass RNA-seq libraries (live-DENV-exposed mosquitoes only, the cohort used to define infection status from carcass viral reads). Row and value conventions are identical to Data S5. Column headers are RNAseq_IDs carrying the suffix ‘_car’ to distinguish carcass libraries from the matched head libraries.

**Data S7. | Differentially expressed genes by comparison, infection defined by DENV reads in the head.** DEGs for every pairwise comparison on the individual-head RNA-seq dataset (DESeq2, Wald test, Benjamini-Hochberg FDR; padj < 0.01, |log₂FC| ≥ 1), with infection status assigned from DENV reads in the head library. One row per gene, with annotation, the full DESeq2 statistics, and the post-hoc outlier-filter output (whether the gene is a clean DEG, carries outlier pools, or is flagged as a likely artifact).

**Data S8. | Immunity and brain gene differential-expression details.** Gene-level differential-expression calls for a curated set of 461 immunity- and brain-specific *Aedes aegypti* genes. Each row is one gene, annotated with its genomic coordinates, functional group, category, and product description, followed by a significant/ns call for every pairwise contrast. DV = live DENV; CN = no-virus control; CH = heat-inactivated DENV control.

**Data S9. | Differentially expressed genes by comparison, infection defined by DENV reads in the carcass.** As Data S7, but with infection status (DVpos/DVneg) assigned from DENV reads in the paired carcass library rather than the head library. One row per gene × comparison; column definitions are identical to Data S7.

**Data S10. | Normalized RNA-seq read counts, tissues from the Neurotranscriptome Atlas**^67^. DESeq2 size-factor-normalized read counts for 14,333 genes across 112 RNA-seq libraries of pooled mosquito tissues. The first column lists *Aedes aegypti* gene identifiers (AaegL5 / VectorBase genome annotation release 68, including the manually curated chemoreceptor genes); each remaining column is one tissue library, identified by its short name. Values are normalized counts (not log-transformed).

## Notes

https://github.com/UmbertoPalatini/DeSeq2_DEGs_OutlierDetector

https://www.ncbi.nlm.nih.gov/bioproject/PRJNA1321547

### Competing Interest Statement

The authors have declared no competing interest.

